# Wnt signaling drives stromal inflammation in inflammatory arthritis

**DOI:** 10.1101/2025.01.06.631510

**Authors:** Alisa A. Mueller, Angela E. Zou, Lucy-Jayne Marsh, Samuel Kemble, Saba Nayar, Gerald F.M. Watts, Cassandra L. Murphy, Emily Taylor, Triin Major, David Gardner, Christopher D. Buckley, Kevin Wei, Soumya Raychaudhuri, Ilya Korsunsky, Andrew Filer, Adam P. Croft, Michael B. Brenner

## Abstract

The concept that fibroblasts are critical mediators of inflammation is an emerging paradigm. In rheumatoid arthritis (RA), they are the main producers of IL-6 as well as a host of other cytokines and chemokines. Their pathologic activation also directly causes cartilage and bone degradation. Yet, therapeutic agents specifically targeting fibroblasts are not available. Here, we find that Wnt receptors and modulators are predominantly expressed in stromal populations in the synovium. Importantly, non-canonical Wnt activation induces robust inflammatory gene expression including an abundance of cytokines and chemokines in synovial fibroblasts *in vitro*. Strikingly, the addition of Wnt ligands or inhibition of Wnt secretion exacerbates or reduces arthritis severity, respectively, *in vivo* in a murine model of inflammatory arthritis. These observations are relevant in human disease, as Wnt activation signatures are enhanced in fibroblasts derived from inflamed RA synovial tissue as well as fibroblasts across other inflammatory diseases. Together, these findings implicate Wnt signaling as a major driver of fibroblast-mediated inflammation and joint pathology. They further suggest that targeting the Wnt pathway is a therapeutically relevant approach to rheumatoid arthritis, particularly in patients who do not respond to conventional treatments and who often express fibroblast-predominant synovial phenotypes.

## Introduction

Rheumatoid arthritis (RA) is characterized by the expansion and inflammatory activation of fibroblasts in the synovium of the joint. These cells, which also support the architecture of the tissue, secrete potent cytokines and other inflammatory molecules that promote leukocyte infiltration and angiogenesis.^1–4^ Moreover, synovial fibroblasts develop an invasive phenotype in RA, forming a destructive pannus that directly mediates bone and cartilage degradation.^1–4^ Recent studies examining synovial biopsy samples from patients refractory to treatment with TNF inhibitors, B cell depletion (rituximab), and IL-6R blockade (tocilizumab) show a fibroblast-enriched transcriptional phenotype. These findings implicate fibroblasts as the major component of treatment-resistant disease.^5–7^

Despite the critical role of fibroblasts in this disease process, the development of therapeutics that specifically target fibroblast populations has remained elusive, and major drivers that potentiate the inflammatory phenotype of fibroblasts is an area of active investigation. Early studies evaluating fibroblast heterogeneity identified key markers defining pathologic and inflammatory fibroblast subsets including Cadherin-11,^8^ Thy-1,^9^ and FAPα.^10^ High-dimensional single cell technologies to molecularly characterize synovial fibroblast subsets have defined 4-10 fibroblast clusters that further define lining and sublining fibroblasts in RA.^7,11,12^ Several fibroblast clusters have been found to be shared across tissues including synovium, lung, gut, and salivary gland including 2 clusters shared across inflammatory diseases in these tissues.^11^ These studies suggest there may be common drivers of pathogenic fibroblast phenotypes across diseases.

The Wnt pathway has been described to play a key role in a number of cellular processes including proliferation and differentiation in various biologic contexts including development and tissue homeostasis.^13–19^ The R-spondin family of proteins have been implicated as activators of Wnt signaling and the Dickkopf family of proteins as inhibitors. Recent reports by the Accelerating Medicines Partnership RA/SLE Network identified a previously unrecognized subset of sublining fibroblasts notable for high expression of Dickkopf-related protein 3 (*DKK3*) and another subset with high expression of R-Spondin 3 (*RSPO3*).^7,12^ Studies of synovium of healthy and hTNFtg mice have suggested that fibroblasts expressing DKK3 may play a role in inflammatory response and matrix remodeling,^20^ and a recent investigation has shown an association of CD200^+^DKK3^+^ fibroblasts with arthritis resolution.^21^ While the molecular functions of *DKK3* are not fully understood, studies have suggested that it can serve as an inhibitor of Wnt signaling and as an immunomodulatory tumor suppressor.^22^ Conversely, *RSPO3* has been shown to augment Wnt signaling and promotes tumor progression.^23^

The discovery of these Wnt-regulated populations led us to hypothesize that the Wnt pathway in the synovium could be acting to regulate the pathogenic behavior of fibroblasts in RA. It is widely assumed that TNF, IL-17, IL-1β and other inflammatory cytokines are the main activators that drive production of IL-6 and other inflammatory factor secretion by fibroblasts. But, the role of Wnt signaling as a major driver of stromal inflammation is a new concept.^24^ Here, we show that Wnt signaling is a major driver of stromal inflammation in rheumatoid arthritis and that targeting this pathway can provide therapeutic benefit. Further, the Wnt activation signature is highly expressed in fibroblasts beyond RA across several autoimmune inflammatory diseases.

## Results

### Wnt pathway members are uniquely expressed in synovial fibroblasts

The Wnt pathway is activated when a Wnt ligand binds to a Frizzled receptor (**Fig. 1a**). Nineteen Wnt ligands and ten Frizzled receptors have been described in humans, and downstream signaling can occur through canonical or non-canonical routes (**Fig. 1a**). There are several modulators of Wnt activity including the R-spondin (RSPO) family of proteins, which enhance Wnt activation, as well as the Dickkopf (DKK) and secreted frizzled-related protein (SFRP) families of proteins that primarily inhibit Wnt signaling.^13,14^

**Figure 1:**
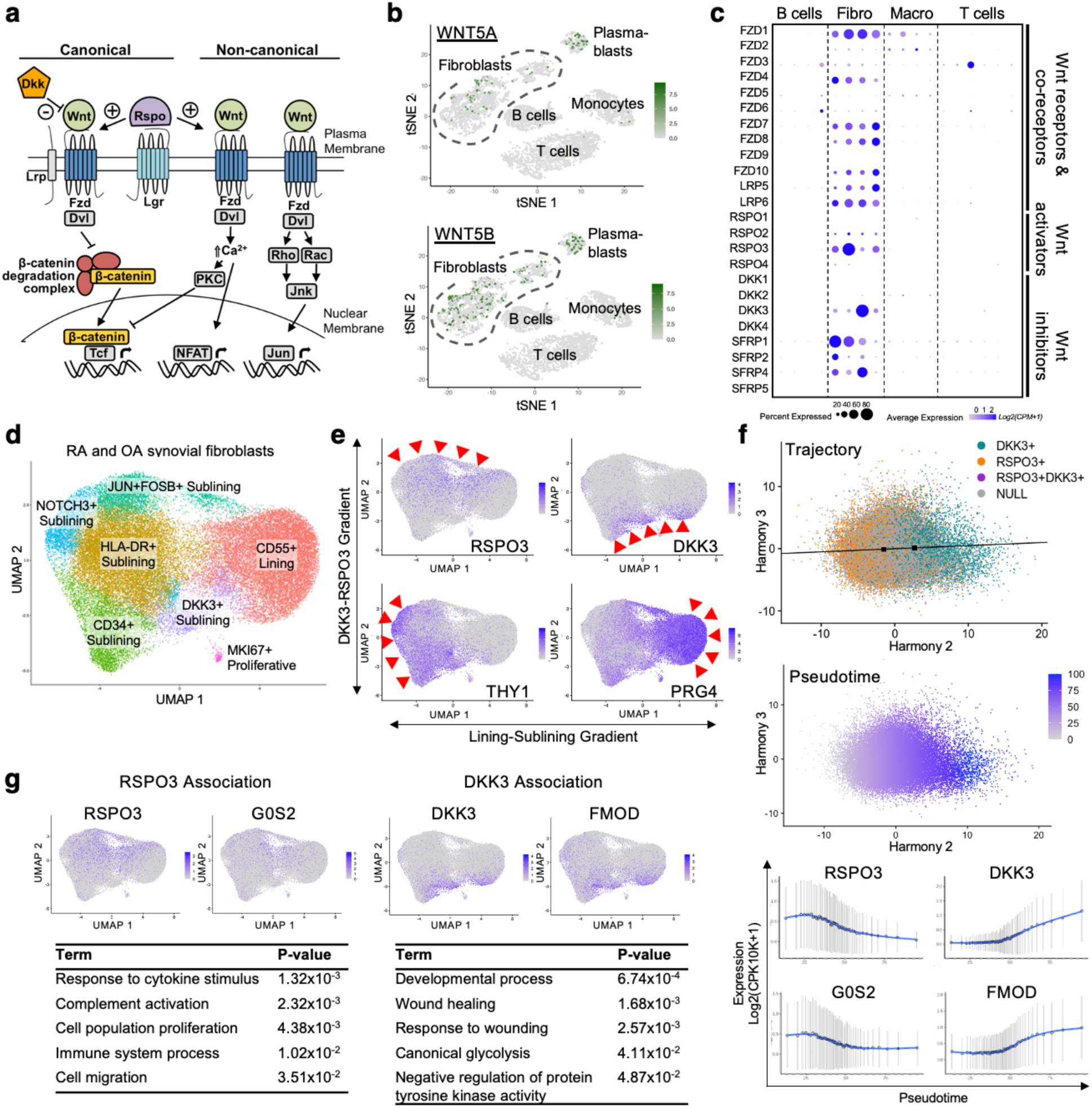
Wnt pathway members are uniquely expressed in synovial fibroblasts, and Wnt regulators DKK3 and RSPO3 form a transcriptional gradient. **a**, Simplified schematic of Wnt signaling illustrating the role of beta-catenin in the canonical pathway as well as calcium signaling and Rho/Rac signaling in the non-canonical pathway. **b**, tSNE plot of scRNA-seq data^12^ from disaggregated and flow-sorted RA and OA synovial tissue. **c**, Dot plot of data from **b** showing expression of Wnt pathway members among synovial populations. Individual clusters are grouped according to the labeled cell type. “Fibro” indicates fibroblasts and “Macro” indicates macrophages. For **b** and **c**, 5,265 sorted cells from n=14 RA and n=3 OA synovial tissue samples are shown. **d**, UMAP depicting fibroblast subpopulations in stromal-enriched scRNA-seq dataset^11^ sorted from RA and OA synovial biopsy samples. **e**, Expression of RSPO3, DKK3, THY1, and PRG4 in UMAP from **d**. Red arrows highlight areas of RSPO3 or DKK3 expression as indicated. **f**, Depiction of trajectory analysis to evaluate RSPO3-DKK3 gradient using a linear trajectory created connecting the centroids of expression of DKK3 and RSPO3 in Harmony-corrected PCA space from dataset in **d** and a pseudotime assigned to the trajectory. The expression of RSPO3 and DKK3 along this trajectory is depicted along with two genes, G0S2 and FMOD, which were identified by hierarchical clustering analysis to show a similar pattern of expression. **g**, Depiction of gene and pathway terms identified to be associated with RSPO3 or DKK3 in the trajectory analysis. For **d-g**, 47,579 cells are projected to UMAP or Harmony-corrected PCA space from OA (n=6) and RA (n=15) samples.

We first sought to determine whether synovial fibroblasts and other synovial cell populations expressed Wnt pathway components. In our analysis of scRNA-seq data from the AMP RA/SLE Network of synovial populations in rheumatoid arthritis and osteoarthritis patients,^12^ we find that several Wnt ligands are expressed throughout synovial populations (**Fig. 1b and Extended Data Fig. 1a and 1b**). However, Wnt receptors, co-receptors and interacting partners that modulate signal transduction within a cell are chiefly expressed by fibroblasts (**Fig. 1c and Extended Data Fig. 1c**). Moreover, of the Wnt ligands represented, most have been associated with non-canonical Wnt signaling including WNT5A, WNT5B, and WNT11. Consistent with this observation, our analyses show that in contrast to canonical Wnt ligands, these non-canonical Wnt ligands do not upregulate AXIN2 in human RA synovial fibroblasts *in vitro* (**Extended Data Fig. 1d**).

### Wnt regulators DKK3 and RSPO3 form a transcriptional gradient that demarcates inflammatory versus remodeling fibroblast phenotypes

In order to fully evaluate the transcriptomic profiles of Wnt signaling pathway members in fibroblasts, we analyzed a stromal-enriched dataset of synovial fibroblasts derived from OA and RA patients (**Fig. 1d and Extended Data Fig. 2**).^11^ In our analysis of this stromal-enriched dataset and that of the AMP/RA SLE Network, we found an unexpected pattern of expression of two Wnt modulators, *RSPO3* and *DKK3*, which play opposing roles in augmenting and putatively inhibiting Wnt signaling, respectively.^15,22^ In particular, single cell gene expression in UMAP projections reveal that *RSPO3* and *DKK3* are expressed in a reciprocal manner (**Fig. 1e**). Moreover, the expression of these genes is not confined to a specific cell cluster. Instead, they form a strong transcriptional gradient of expression that is present across both lining and sublining fibroblasts and is expressed in a distinct pattern from the lining-sublining (PRG4-THY1) fibroblast differentiation gradient facilitated by the Notch signaling pathway (**Fig. 1e**). Histological multispectral immunofluorescent staining data of RA synovial tissue confirms reciprocal expression of *RSPO3* and *DKK3* (**Extended Data Fig. 3**).

To explore whether the *RSPO3*-*DKK3* gradient was correlated with a transition in fibroblast phenotype, we sought to identify co-varying genes that follow a similar expression gradient to *RSPO3* and *DKK3*. To this end, we constructed a linear trajectory along the *RSPO3*-*DKK3* gradient and assigned each cell a pseudotime value based on its position along this trajectory. We then used a hierarchical clustering approach to identify genes that follow the same patterns of expression as *RSPO3* and *DKK3* (**Fig. 1f**). For example, G0/G1 switch gene 2 (G0S2), a regulator of metabolism and proliferation, and fibromodulin (FMOD), a gene involved in collagen assembly, show similar patterns of expression to RSPO3 and DKK3, respectively (**Fig. 1f**). Our analyses showed that cells expressing *RSPO3* are associated with genes enriched for response to cytokine stimulus, complement activation, proliferation, and migration among others suggesting an activated, inflammatory phenotype (**Fig 1g**). By contrast, genes that followed a similar expression pattern to *DKK3* were enriched for gene ontology terms including developmental process, wound healing, glycolysis, and inhibition of receptor tyrosine kinase signaling, which pointed to a tissue remodeling phenotype (**Fig. 1g**).

### Wnt signaling promotes an inflammatory phenotype of synovial fibroblasts

In light of our data demonstrating that synovial fibroblasts are poised to signal through the Wnt pathway and the potential relevance of the Wnt transcriptional gradient to cell phenotype, we sought to comprehensively evaluate the impact of Wnt activation on synovial fibroblast gene expression. To this end, we stimulated RA synovial fibroblast cell lines with Wnt ligands across a concentration range and evaluated the transcriptional response at two time points (**Fig. 2a**). Wnt-3a and Wnt-5a, which represent prototypical canonical and non-canonical Wnt ligands, respectively, were applied.

**Figure 2:**
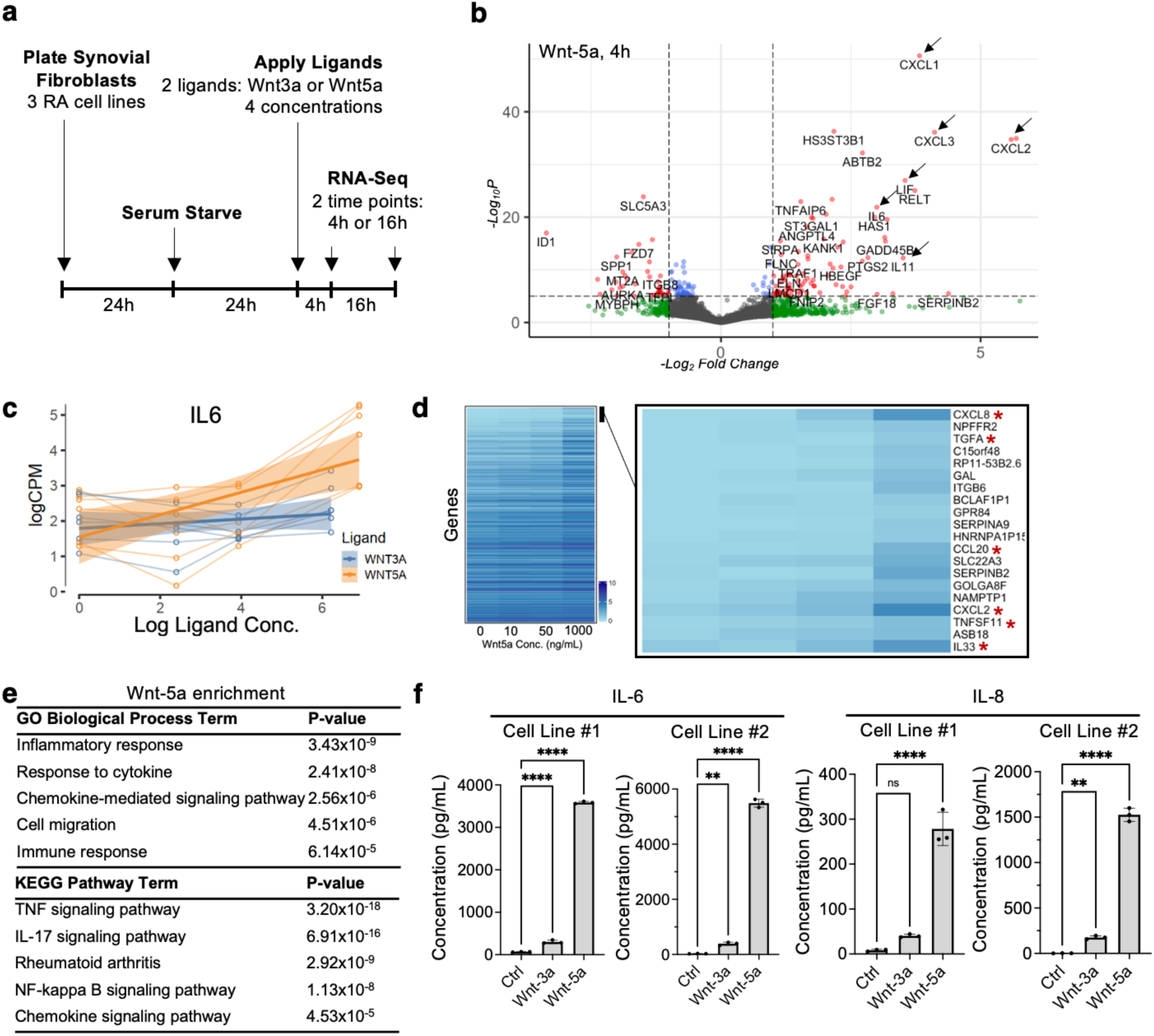
Wnt signaling promotes an inflammatory phenotype of synovial fibroblasts. **a**, Schematic depicting parameters of *in vitro* stimulation of synovial fibroblasts with Wnt ligands. **b**, Volcano plot depicting change in gene expression of RA synovial fibroblasts treated with Wnt 5a ligand for 4 hours (n=3 biologic replicates per condition). **c**, Graph showing change in IL6 expression with increasing concentrations of Wnt-3a or Wnt-5a. **d**, Heatmap showing expression of individual genes (rows) with increasing Wnt-5a ligand concentration (columns). Genes are ordered by the degree to which their expression increases in response to increasing concentrations of Wnt-5a. The genes with the highest increase in expression are shown to the right. Asterisks indicate inflammatory mediators. For **c** and **d**, n=3 biological replicates per condition. **e**, Selected GO Terms and KEGG pathway terms that show enrichment in genes upregulated by Wnt-5a stimulation. **f**, ELISA depicting IL-6 and IL-8 secretion by RA synovial fibroblasts in response to stimulation with Wnt-5a, Wnt-3a, or a PBS control, where n=3 for each condition. Cell lines #1 and #2 denote two individual RA synovial fibroblast cell lines. Mean ± SD shown. To calculate significance, an ordinary one-way Anova with Holm Šídák’s multiple comparisons test was used. **p<0. 001, ****p<0.0001.

Strikingly, Wnt stimulation induced a profound upregulation of genes encoding inflammatory cytokines and chemokines. For instance, differential gene expression analyses showed pro-inflammatory cytokines including *IL6*, *LIF*, *CXCL1*, *CXCL2*, *CXCL3*, and *IL11* and others were among the most highly upregulated genes after stimulation with Wnt-5a for 4 hours (**Fig. 2b**). Multivariate regression analyses were also used to identify genes that show a dose-responsive increase in expression with increasing Wnt ligand concentration. As an example, the expression of *IL6* is shown to increase with increasing Wnt-5a concentration (**Fig. 2c**). The expression of all genes across Wnt-5a concentrations were fitted to a line and the slope calculated in order to rank genes with those showing the largest positive correlation (higher slope) ranked highest (**Fig. 2d**). In agreement with the results for differential gene expression, several inflammatory mediators including *CXCL8*, *TGFA*, *CCL20*, *CXCL2*, *TNFSF11*, and *IL33* were among the genes with the highest positive correlation to increasing Wnt-5a dose (**Fig. 2d**). Additionally, gene set enrichment analyses of the genes upregulated by Wnt-5a stimulation identified pathways known to be relevant to RA inflammatory pathology including TNF signaling pathway, IL-17 signaling pathway, and NF-kappa B signaling pathway among others (**Fig. 2e**). Our studies also showed enhanced cell proliferation in response to Wnt-5a (**Extended Data Fig. 4**).

Of note, while both canonical (Wnt-3a) and non-canonical (Wnt-5a) activation resulted in upregulation of inflammatory cytokines, we found that non-canonical Wnt activation resulted in a more robust induction of the inflammatory fibroblast phenotype. For example, while IL6 transcript levels are increased with Wnt-5a stimulation, it is only modestly upregulated with Wnt-3a stimulation (**Fig. 2c**). Increases in secreted IL-6 and IL-8 levels in response to non-canonical Wnt-5a stimulation were confirmed, and Wnt-5a was found to upregulate IL-6 and IL-8 levels more robustly than canonical Wnt ligand Wnt-3a (**Fig. 2f**).

### Wnt-5a activation and RSPO3 enrichment scores are associated with inflammatory disease

Given the predominance of non-canonical Wnt ligands in the synovium and the profound effect of non-canonical Wnt signaling on the production of inflammatory mediators, we endeavored to determine whether Wnt activation had clinical relevance to RA. To address this question, we developed a Wnt-5a activation signature composed of genes that were upregulated upon Wnt-5a stimulation. We first analyzed an scRNA-seq dataset^11^ enriched for stromal fibroblast cell populations in synovial samples from patients with RA and OA controls as well as other inflammatory diseases. We used a weighted scoring system to calculate a Wnt-5a activation score for each fibroblast cell based on the level of expression of the 50 most upregulated Wnt-5a-induced genes. Expression of the Wnt agonist, *RSPO3*, is associated with a high Wnt-5a activation score, suggesting higher Wnt activation in *RSPO3*-expressing cells (**Fig. 3a**). Conversely, *DKK3*-expressing fibroblasts exhibit low Wnt-5a activation scores (**Fig. 3a**).

**Figure 3:**
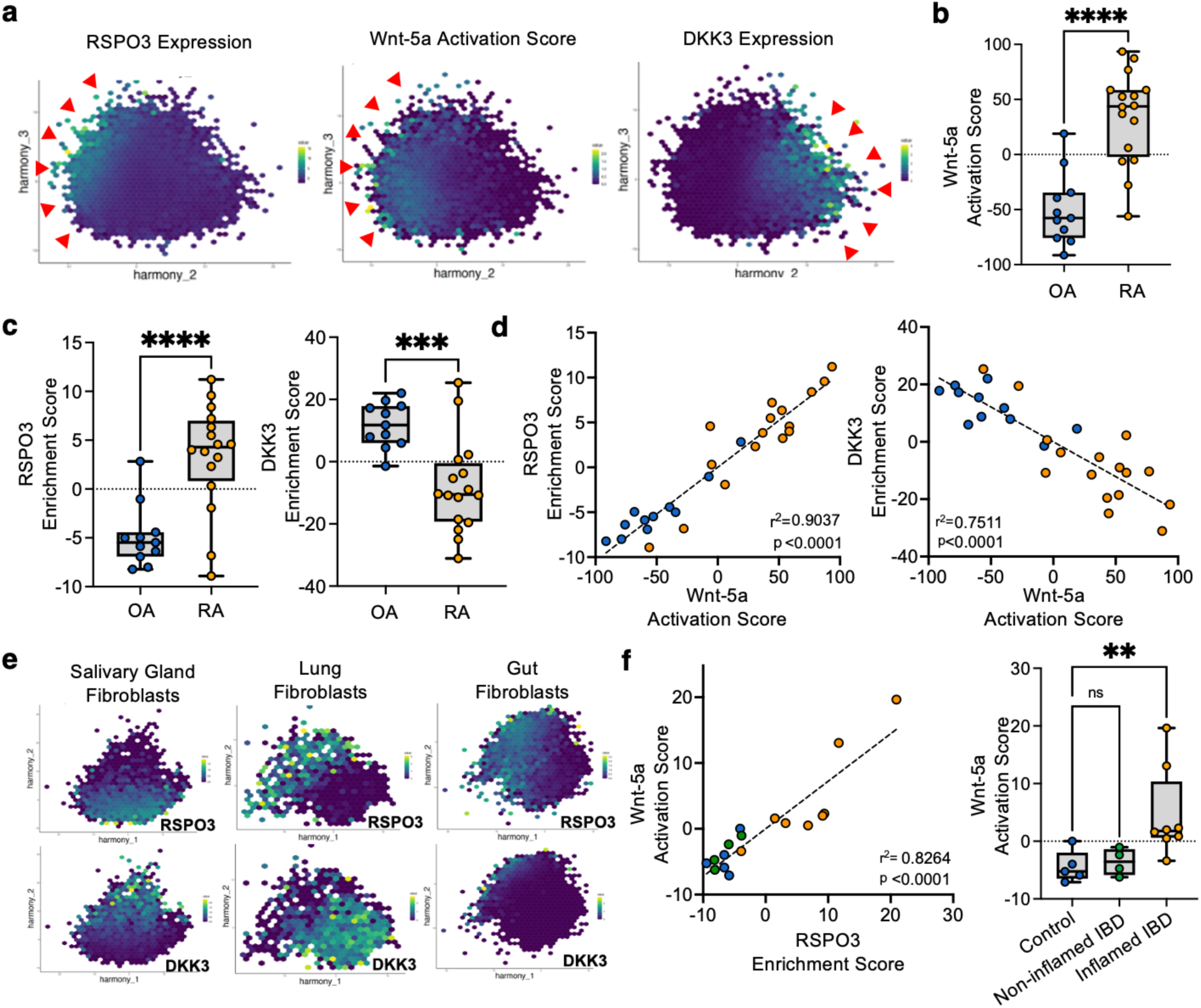
Enhanced Wnt signaling defined by RSPO3-DKK3 expression is associated with stromal inflammation in rheumatoid arthritis synovium and more broadly in other tissues. **a**, Harmony-corrected PCA plot of scRNA-seq data^11^ from 47,812 OA and RA synovial fibroblasts are shown with Wnt-5a activation scores, RSPO3 expression, and DKK3 expression depicted. Red arrows indicate areas of increased expression. **b**, Wnt-5a activation scores are shown for OA (n=11) and leukocyte-rich RA (n=16) synovial samples from bulk RNA sequencing data.^12^ **c**, RSPO3 and DKK3 enrichment scores are shown for OA (n=11) and leukocyte-rich RA (n=16) synovial samples from bulk RNA sequencing data collected by the Accelerated Medicines Partnership.^12^ **d**, Graph depicting correlation of Wnt-5a activation scores with RSPO3 (left) and DKK3 (right) enrichment scores for bulk RNA sequencing data of OA (n=11) and leukocyte-rich RA (n=16) synovial fibroblasts.^12^ **e**, Harmony-corrected PCA plot depicting RSPO3 and DKK3 expression in fibroblasts of the salivary gland from patients with Sjogren’s disease (n=7) or sicca symptoms (n=6), lung tissue from patients with ILD (n=19) or control (n=4), and colon from patients with inflammatory bowel disease (IBD) (n=8) or control (n=5) from samples collected by the Roche Fibroblast Network Consortium.^11^ **f**, On the left, a graph of Wnt-5a activation scores and RSPO3 enrichment scores are shown for gut fibroblast samples from healthy control patients and patients with IBD. On the right, Wnt-5a enrichment scores are shown for synovial fibroblast samples obtained from healthy control patients (n=5), non-inflamed IBD (n=4) and inflamed IBD (n=8).

We then analyzed bulk RNA sequencing data of synovial fibroblasts sorted from leukocyte-rich RA synovial samples and OA control synovial samples.^12^ Using the same scoring system, we calculated a Wnt-5a activation score for each patient sample based on the level of expression of the top 50 genes most highly upregulated by Wnt-5a. We also calculated *RSPO3* and *DKK3* enrichment scores using genes that were identified through covarying genes identified by trajectory analyses shown in **Fig. 1f and 1g**. Notably, we found that the Wnt-5a activation score is enriched in synovial fibroblasts in RA compared to OA (**Fig. 3b**). Moreover, the *RSPO3* enrichment scores were higher in RA compared to OA with a striking positive correlation to Wnt-5a activation scores (**Fig. 3c and 3d**). Conversely, the *DKK3* enrichment score was decreased in RA with a negative correlation with Wnt-5a activation scores (**Fig. 3c and 3d**).

In the setting of evidence for strong Wnt activation in RA, we explored whether other chronic inflammatory diseases would show similar patterns of Wnt expression and activation in fibroblasts. We analyzed publicly available data of fibroblasts from the following disease subsets: salivary gland fibroblasts in Sjogren’s disease vs. patients with non-autoimmune sicca symptoms; lung tissue from interstitial lung disease vs. controls; and inflamed intestinal tissue from ulcerative colitis and healthy controls.^11^ In these tissues RSPO3 and DKK3 show distinct reciprocal expression patterns in Sjogren’s disease and in interstitial lung disease (**Fig. 3e**). While the reciprocal expression is not present in the gut fibroblasts (**Fig. 3e**), the fundamental connection of Wnt to inflammation is preserved as there is a positive association of Wnt-5a activation scores and *RSPO3* enrichment signatures. Moreover, there is an upregulation of Wnt-5a activation scores in fibroblasts from inflamed intestine (**Fig. 3f**).

### Intra-articular activation of non-canonical Wnt signaling worsens inflammatory arthritis

Given the potential importance of Wnt signaling in rheumatoid arthritis and other inflammatory conditions, we tested whether modulation of Wnt signaling could impact arthritis outcomes in mouse models of inflammatory arthritis. First, we evaluated whether Wnt signaling patterns in mouse models resembled those seen in human. In the K/BxN serum-transfer arthritis mouse model, serum from transgenic, arthritic K/BxN mice is transferred to naïve mice (**Extended Data Fig. 5a**). We evaluated scRNA-seq data of synovial fibroblasts isolated from the digested synovia of murine joints at time points representing, resting (day 0), peak arthritis (day 9), resolving arthritis (day 15), and fully resolved arthritis (day 22) (**Extended Data Fig. 5b and 5c**). We find that expression patterns of Wnt pathway components, including the reciprocal expression RSPO3 and DKK3, mimic those seen in human patients (**Extended Data Fig. 5d**). Moreover, in the setting of our data supporting non-canonical Wnt pathway enrichment in RA, we proposed that Wnt-5a signatures would be upregulated at peak arthritis. Indeed, both Wnt-5a activation and RSPO3 trajectory signatures are enriched in synovial fibroblasts collected at the peak of arthritis (day 9) compared to control and decreases upon the resolution of arthritis (day 22) (**Fig. 4a**).

**Figure 4:**
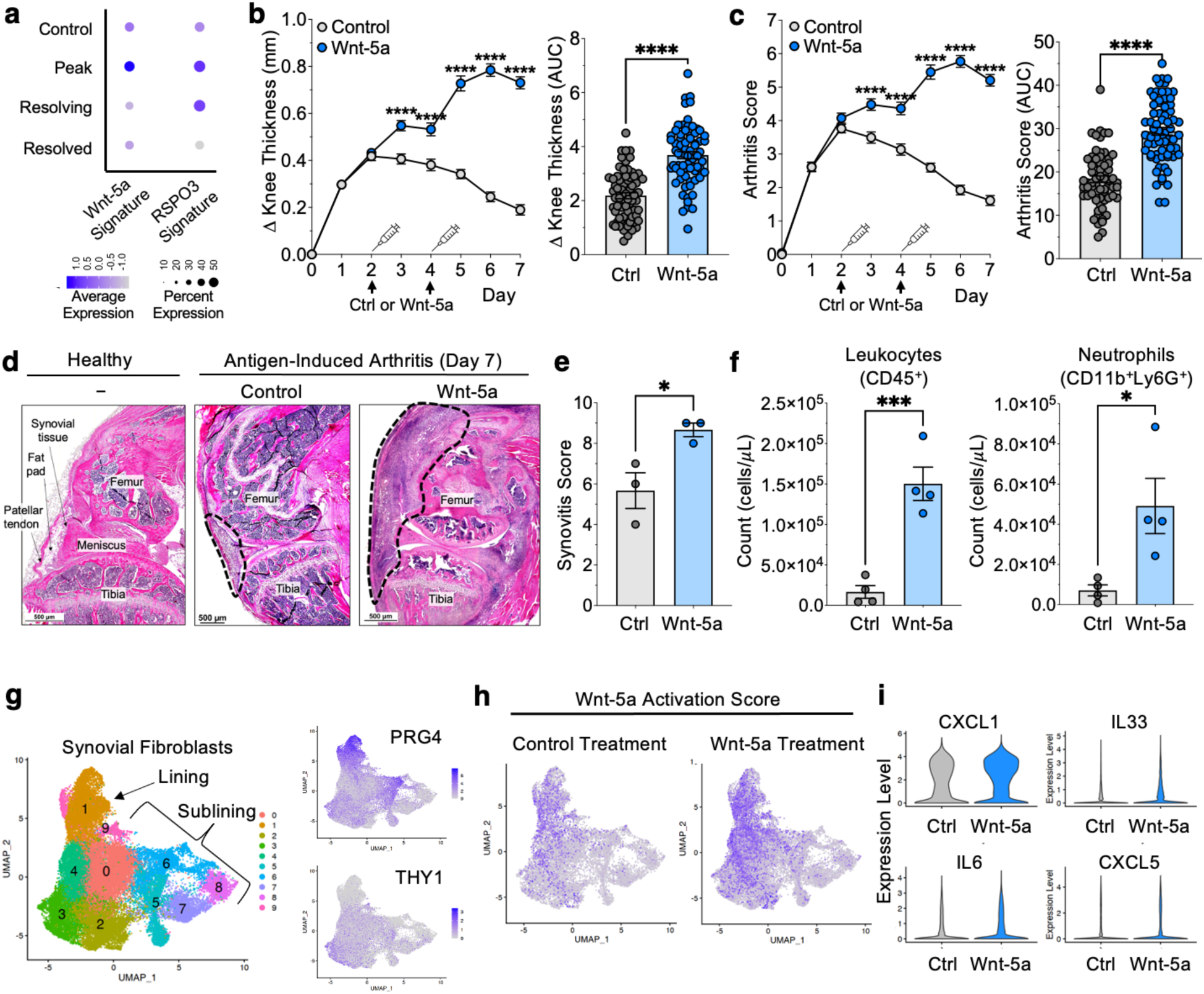
Stimulation of Wnt signaling increases inflammation and arthritis severity in a murine model of inflammatory arthritis. **a**, Dot plot showing enrichment of RSPO3 trajectory signatures and Wnt-5a enrichment signatures across indicated time points in synovial fibroblasts in K/BxN inflammatory arthritis model (n=3 biologic replicates per time point). **b,c**, Knee thickness measurements (**b**) and global arthritis scores (**c**) of mice undergoing AIA that received intraarticular injection with recombinant Wnt-5a (n=67) or control (n=69) at the indicated time points. For **b** and **c**, mean ± SEM is shown. For the time courses in **b** and **c**, significance at each time point was determined by a repeated measures two-way ANOVA with Šídák’s multiple comparisons test. For the AUC bar plots, normality assessed with the Shapiro Wilk test and a Student’s T test was used. **d**, Histology depicting knee anatomy in a healthy mouse (no AIA) or in mice that underwent AIA and were treated with recombinant Wnt-5a or a PBS control. Dashed lines outline synovial infiltrate. **e**, Synovitis scores of knee synovial samples by histology in mice treated with control (n=3) or Wnt-5a (n=3). Mean ± SEM is depicted. A Student’s t test was used to determine significance. **f**, Flow cytometry analysis of knee synovial tissue from Wnt-5a-treated mice (n=4) and controls (n=4) showing absolute cell counts for leukocytes, and neutrophils. Normality was assessed with the Shapiro Wilk test, and the Student’s T test was applied. **g**, UMAP depicting scRNA-seq data for synovial fibroblasts isolated from knee joints of Wnt-5a- and control-treated mice. Expression of lining marker PRG4 and sublining marker THY1 are depicted. **h**, UMAP showing Wnt-5a activation scores from scRNA-seq of knee-derived synovial fibroblasts isolated from Wnt-5a-treated mice (n=3) or PBS-treated controls (n=3). **i**, Violin plots showing expression of CXCL1, IL33, IL6, and CXCL5 for mice treated with control PBS (n=3) or Wnt-5a (n=3). *p<0.05, ***p<0.001, ****p<0.0001. AUC, area under the curve; Abs, absolute.

Since non-canonical Wnt stimulation induced an inflammatory fibroblast phenotype and Wnt signatures were enriched in human RA and at the peak of arthritis in mice, we predicted that enhancing non-canonical Wnt activation would heighten arthritis severity. We tested this hypothesis in an antigen-induced arthritis (AIA) model where mice were primed with methylated BSA (mBSA) at day −21 prior to the induction of arthritis by local injection of mBSA into the knee joint. This model allowed us to localize the effects of Wnt-5a by also injecting it directly into the knee joint of C57Bl/6 mice on days 2 and 4 after the mBSA joint injection. Intriguingly, injection of Wnt-5a increases the amplitude and duration of knee swelling (**Fig. 4b**) and also led to an increase in global arthritis scores (**Fig. 4c**). This worsening clinical phenotype was associated with increased lining hyperplasia and leukocyte infiltration as observed by histology (**Fig. 4d and 4e**). Flow cytometry studies of synovial tissues showed a significant increase in leukocytes, particularly neutrophils and macrophages among others (**Fig. 4f and Extended Data Fig. 6**).

While absolute counts of fibroblast populations did not differ (**Extended Data Fig. 6c**), transcriptomic analyses demonstrated inflammatory activation of these cells. In particular, there were enhanced Wnt enrichment signatures in sublining and lining fibroblasts where Wnt-5a was injected versus vehicle control (**Figs. 4g and 4h**), and an increased expression of cytokines and other inflammatory molecules was observed (**Fig. 4i**).

### Inhibition of Wnt signaling ameliorates arthritis severity

Next, we evaluated whether inhibition of Wnt signaling could decrease inflammatory arthritis severity. We utilized the AIA model and treated mice with LGK974, a chemical inhibitor of Wnt ligand secretion that prevents signaling through non-canonical and canonical signaling pathways.^25^ We used systemic administration of LGK974 since it was injected multiple times during the course of arthritis. It abrogated arthritis by reducing knee swelling (**Fig. 5a**) and global arthritis scores (**Fig. 5b**). A subset of mice that underwent AIA-induced arthritis for 7 days showed a persistent decrease in knee swelling in LGK974-treated mice (**Extended Data Fig. 7**). Tissues were collected for additional studies at Day 4 when knee swelling was decreased maximally in LGK974-treated mice compared to vehicle-injected control mice. Decreased lining hyperplasia and leukocyte infiltration was observed in histology of LGK974-treated mice compared to control (**Fig. 5c and 5d**). In addition, flow cytometry analyses revealed decreased leukocyte numbers and a trend toward decreased neutrophils (**Fig. 5e**). Similar to the case with Wnt administration, synovial fibroblast numbers were unchanged (**Extended Data Fig. 8**). However, scRNA-seq analyses of murine synovial fibroblasts (**Fig. 5f**) demonstrated that administration of LGK974 was associated with a decrease in CXCL5 (**Fig. 5g**), a chemokine known to recruit neutrophils in RA^26^ and which was increased with Wnt activation. Moreover, fibroblasts showed upregulation of inhibitors of JAK signaling, namely SOCS1 and SOCS3 (**Fig. 5h**).^27^ Together, these findings suggest a potential mechanistic explanation for reduced inflammation and arthritis severity in mice undergoing Wnt inhibition.

**Figure 5:**
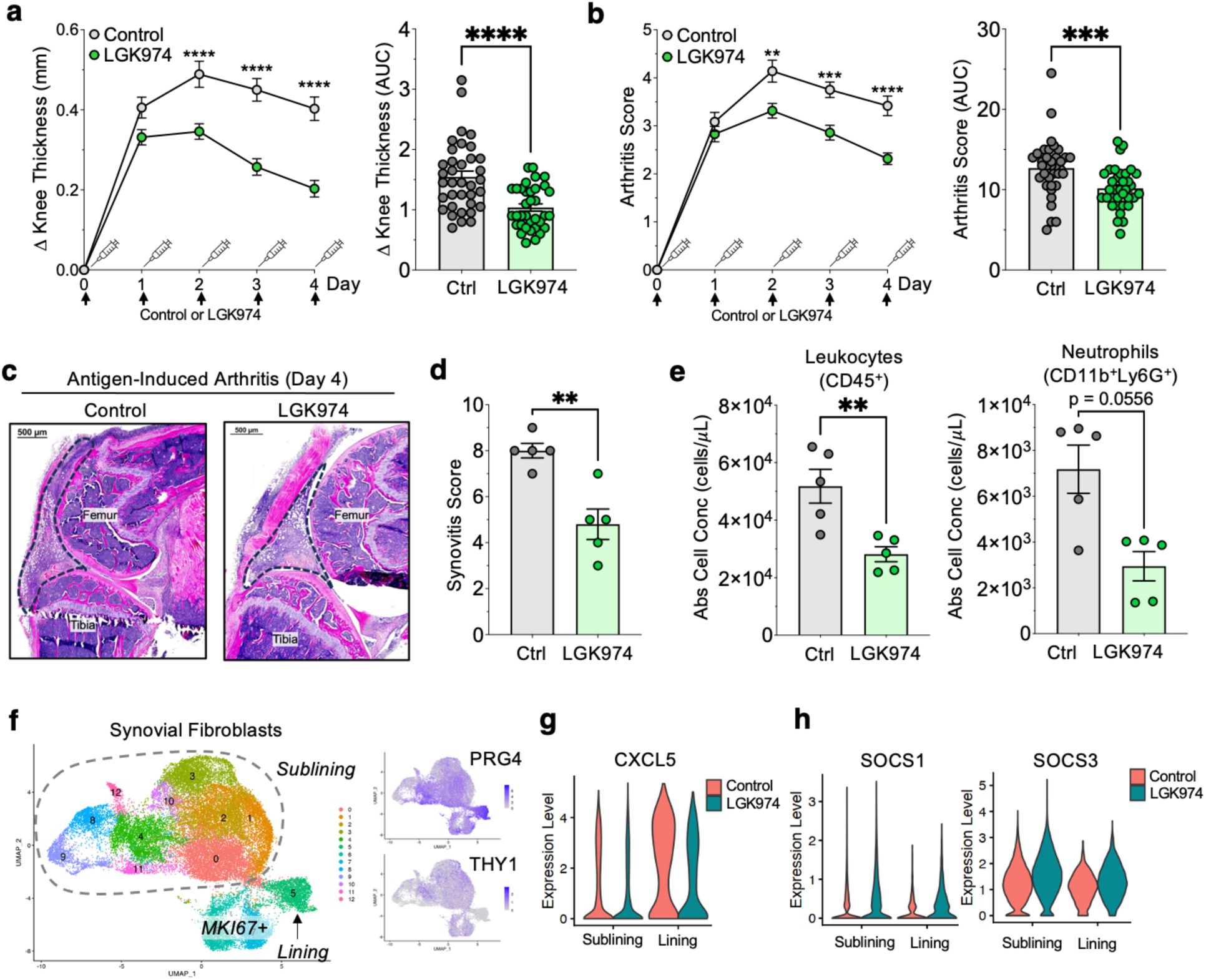
Inhibition of Wnt signaling decreases inflammatory infiltrate and reduces arthritis severity. **a-b**, Knee thickness measurements (**a**) and global arthritis scores (**b**) of mice undergoing AIA that were treated with recombinant LGK974 (n=35) or a vehicle control (n=36) at the indicated time points. **c**, Representative H&E of knee joints in mice that underwent AIA and were treated with recombinant LGK974 or a vehicle control. **d**, Synovitis scores of mouse knee synovial samples in vehicle control- (n=5) and LGK974- (n=5) treated mice based on histologic parameters as described in the Methods. **e**, Flow cytometry analysis showing absolute cell counts for leukocytes, neutrophils, and synovial macrophages in LGK974 treated (n=5) or control mice (n=5). **f**, UMAP of 34,728 showing synovial fibroblasts harvested from knee joints of LGK974- (n=3) or control-treated (n=3) mice with the expression of lining marker PRG4, sublining marker THY1, and proliferation marker MKI67 denoting major populations. **g,h**, Violin plots depicting expression of CXCL5, SOCS1, and SOCS3 in synovial fibroblasts from sublining and lining populations in LGK974 or control-treated mice. For **a**, **b**, **d**, and **e**, mean ± SEM is depicted. For the time courses shown in **a** and **b**, significance at each time point was determined by a repeated measures two-way ANOVA with Šídák’s multiple comparisons test. For the AUC bar plots, normality assessed with the Shapiro Wilk test and a Student’s T test was used for **a** and Mann-Whitney test for **b**. For **d** and **e**, normality was assessed with the Shapiro Wilk test, and the Student’s T test was applied to **d** and the leukocyte bar plot in **e**. The Mann-Whitney test was used for the neutrophil bar plot in **e**. *p<0.05, **p<0.01, ***p<0.001, ****p<0.0001. AUC, area under the curve; Abs, absolute.

## Discussion

Here, we demonstrate that Wnt pathway members are predominantly expressed in stromal populations. Importantly, non-canonical Wnt signaling drives the inflammatory activation of synovial fibroblasts, and synovial fibroblasts in RA show enrichment in transcriptional signatures of Wnt activation. Finally, modulation of this pathway in a murine model of inflammatory arthritis determines arthritis severity.

Prior studies have implicated various cytokines in RA pathogenesis. These cytokines include TNF, IL-17, leukemia inhibitory factor (LIF), and IL-1β, which can act independently and synergistically to promote inflammatory cytokine secretion in synovial fibroblasts.^28–31^ Furthermore, IFNγ^32^ and cell adhesion molecules, like Cadherin-11,^33^ have been demonstrated to mediate synovial fibroblast invasiveness.^8,33^ Recently, Notch signaling has been shown to promote differentiation of synovial fibroblasts into inflammatory sublining subtypes.^34^ Unexpectedly, our investigation demonstrates that Wnt signaling serves as an independent mechanism that is a major regulator of inflammatory fibroblast activation in RA.

Historically, the Wnt pathway has been prominently associated with its roles in processes independent of inflammation including cell proliferation, differentiation, and migration in contexts including tissue homeostasis, fibrosis, and cancer.^14,15^ However, recent studies have begun to point to its functions in regulating tissue inflammation and immune cell responses in autoimmunity and cancer. For example, one study showed that the activation of β-catenin in T cells of mice facilitated intestinal and colonic inflammation. Intriguingly, reports have also shown an immune suppressive function of Wnt.^35^ Many studies have focused on the role of canonical Wnt signaling in modulation of leukocyte biology, while the impact on fibroblast biology is less clear.^24,36^ In our investigation, we demonstrate the importance of Wnt signaling in fibroblast inflammatory pathogenicity. Moreover, while both canonical and non-canonical Wnt signaling alter fibroblast phenotype, it is the activation of the non-canonical pathway that had a striking impact on promoting the inflammatory phenotype of synovial fibroblasts, which is linked to RA and other inflammatory diseases.

In the context of RA, there are limited studies that have evaluated the impact of the Wnt pathway on disease biology. In synovial tissue and fibroblasts isolated from RA patients, *WNT5A* expression was enriched,^37^ and expression of Wnt1 inducible signaling pathway protein-3 was found to be elevated compared to expression in fibroblasts from healthy and OA patients.^38^ Transfection of normal synovial fibroblasts cells with a *WNT5A* expression vector increased transcriptional expression of IL-6, IL-8, and IL-15,^37^ and transfection with a Wnt-1 expression vector promoted proMMP-3 expression.^39^ However, these studies were performed *in vitro* and focused on factor production but did not examine the effect of Wnt on transcriptomic signatures and cell states or in animal models of arthritis. Importantly, in our study, we show through comprehensive analysis of Wnt stimulation at sequential doses and time points that Wnt activation leads to upregulation of IL6, LIF, CXCL1, CXCL2, and CXCL3 among other genes. These cytokines and chemokines are known to be critical in RA pathogenesis by promoting inflammatory potentiation of leukocytes and stromal cells, immune cell recruitment, and cartilage and bone destruction.^3^ We use this transcriptome-wide analysis to generate Wnt activation signatures and show an enrichment in RA as well as demonstrate their roles in animal models of arthritis.

Moreover, our analyses of human single cell data from synovial fibroblasts revealed surprising patterns of Wnt pathway-related transcriptional profiles. On one hand, we identify individual synovial fibroblast clusters or cell states characterized by high *DKK3* or *RSPO3* expression akin to prior studies.^7,12^ However, our data show that the enrichment of Wnt signatures follow a transcriptional gradient that is not best defined by particular clusters, but instead runs across clusters in a more global fibroblast gradient. Interestingly, the non-canonical Wnt activation signature was positively correlated with expression of the Wnt activator, *RSPO3*, and negatively correlated with the putative Wnt antagonist, *DKK3*. This gradient is independent of other transcriptional gradients recently described including the Notch signaling gradient.^34^ These findings suggest a unique role for this pathway in regulating fibroblast biology across anatomic and cell cluster designations.

In another context, the Wnt pathway has been implicated in osteoarthritis where its activity has been associated with chondrogenesis and osteogenesis.^40^ In a mouse model of meniscal injury, canonical Wnt inhibition with small-molecule inhibitor XAV-939 decreased cartilage degeneration and OA severity.^41^ Likewise, the Wnt antagonist sclerostin has been associated with chondrocyte function, and its modulation affects OA severity.^42^ Additionally, the small molecule inhibitor lorecivivint (SM04690), which targets intranuclear kinases downstream of the Wnt pathway, has shown promise in ongoing clinical trials.^43,44^

Prior animal studies addressing the role of the Wnt pathway in inflammatory arthritis have primarily focused on bone remodeling.^45–47^ *Wnt5a* has been shown to impact osteoclast activity in a murine model of inflammatory arthritis where its deletion in a K/BxN serum transfer model led to a reduction in cartilage degradation and decrease in osteoclast activity.^46^ Conversely, in a separate study, deletion of *Wnt9a* was shown to worsen arthritis severity in a transgenic TNF mouse model without major impact on arthritis outcomes K/BxN serum transfer mice.^48^ Though given the minimal expression of *WNT9A* in human synovium, the relevance of this model is unclear. In both studies, the impact of Wnt modulation in fibroblasts and potential for a pharmacologic approach to impact inflammatory arthritis outcomes is not known.

In our study, we leverage transcriptomic analyses to show the relevance of the Wnt pathway in the time course of arthritis flare as well as the importance of this pathway in determining the outcome of joint inflammation in mouse models of arthritis. The use of LGK974 to abrogate Wnt signaling was particularly appealing for our study as it has precedence for use in multiple animal models^25,49,50^ and has been tested in human clinical trials for malignancy where phase I studies have shown reasonable safety.^51^ Here, we demonstrate the impact of Wnt modulation on leukocyte infiltration through histology and flow cytometry. Furthermore, scRNA-seq analyses of the mouse synovium show the impact of Wnt modulation on inflammatory fibroblast phenotype *in vivo*.

These experiments have particular clinical relevance to RA where TNF inadequate responders represent an important unmet need, and it is estimated that less than 50% of patients with RA are in remission.^52^ Moreover, 10-25% of patients are refractory to treatment with all conventional disease modifying antirheumatic drugs and biologics.^53,54^ Intriguingly, comprehensive analyses of synovial tissue biopsies from RA patients suggests that RA treatment non-responders show decreased leukocyte infiltration and demonstrate a fibroblast-enriched tissue pathotype.^55,56^ These studies suggest that synovial fibroblasts may play a critical role in treatment-refractory RA. Thus, targeting fibroblasts therapeutically is an area of compelling interest.

In our analyses we demonstrate that human synovial fibroblasts from RA patients show marked enrichment of Wnt activation signatures, and this is a pattern shared across other inflammatory diseases. Our work indicates that Wnt signaling may be an important driver of inflammation across several diseases and that targeting the Wnt pathway may be a compelling mechanism, particularly in patients that are refractory to other treatments.

## Acknowledgements

A.A.M. received funding from NIH NIAMS K08AR083513, the Rheumatology Research Foundation Tobé and Stephen E. Malawista, MD, Endowment in Academic Rheumatology, NIH NIAMS T32AR007530, the Joint Biology Consortium Microgrant P30AR070253. A.Z. received funding from NIH NIAID F30 AI174699 from NIH NIGMS T32GM007753 and T32GM144273. K.W. Is supported by a NIH NIAMS K08AR077037, Rheumatology Research Foundation Innovative Research Award, Burroughs Wellcome Fund Career Award in Medical Sciences, and Doris Duke Foundation Clinical Scientist Development Award. S.R. received funding from NIH U01 HG012009, NIH R01 AR063709, NIH P01 AI148102, and NIH UC2 AR081023. A.P.C. is supported by a Kennedy Trust for Rheumatology Research Senior Fellowship [KENN192006], Versus Arthritis Endowment [17553] and Foundation for Rheumatology Career grant [project 038]. M.B.B. received funding from NIH R01AR0637039 and NIH P01AI148102. We thank Dr. Harris Perlman for contribution of K/BxN serum for serum transfer arthritis experiments.

## Competing Interests

K.W. is consultant for Mestag Therapeutics and Gilead Sciences and reports grant support from Gilead Sciences, Merck, and 10X Genomics. S.R. is a founder for Mestag, Inc. and also serves as a scientific advisor to Sonoma, Ground Truth, Janssen, Pfizer, Nimbus, and Third Rock Ventures. I.K. is a consultant for Mestag Therapeutics. A.F. has provided consultancy for Johnson & Johnson and Sonoma and has received institutional grant funding from BMS, Roche, UCB, Nascient, Mestag, GSK, Johnson & Johnson and Synact. M.B.B. is a consultant to GSK, Moderna, and 4F0 Ventures and is a founder of Mestag Therapeutics.

---

**Extended Data Figure 1:**
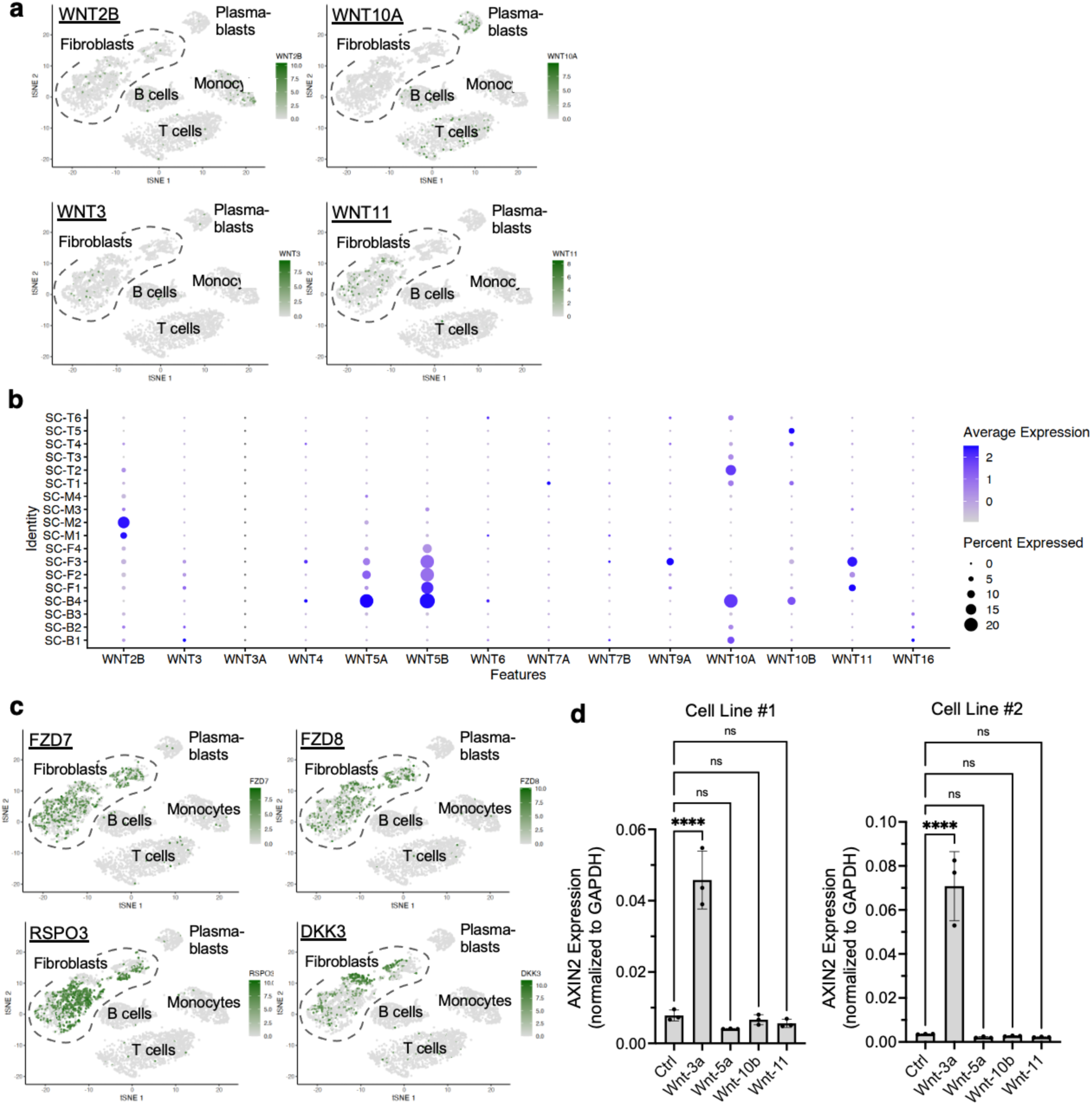
Expression of Wnt pathway members in human RA and OA synovial fibroblasts. **a**, tSNE plot showing expression of the indicated Wnt ligands from of scRNA-seq data^12^ of RA and OA synovial tissue. **b**, Dot plot showing expression of Wnt ligands among synovial populations using the dataset from (**a**). B1-B4 are B cell subsets, F1-F4 are fibroblast subsets, M1-M4 are monocyte subsets, and T1-T6 are T cell subsets. **c**, tSNE plot showing scRNA0seq from dataset from (**a**) showing expression of FZD7, FZD8, RSPO3, and DKK3. **d**, Bar graph of qPCR showing expression of AXIN2 in response to stimulation with the indicated Wnt ligands. STB-020 and RA190321 are individual RA cell lines (n=3 well replicates per condition). Mean ± SD is depicted. Significance was determined using a one-way ANOVA with Dunnett’s multiple comparisons test ****p<0.0001. In A-C, 5,265 sorted cells are shown with n=3 OA samples and n=14 RA samples.

**Extended Data Figure 2.**
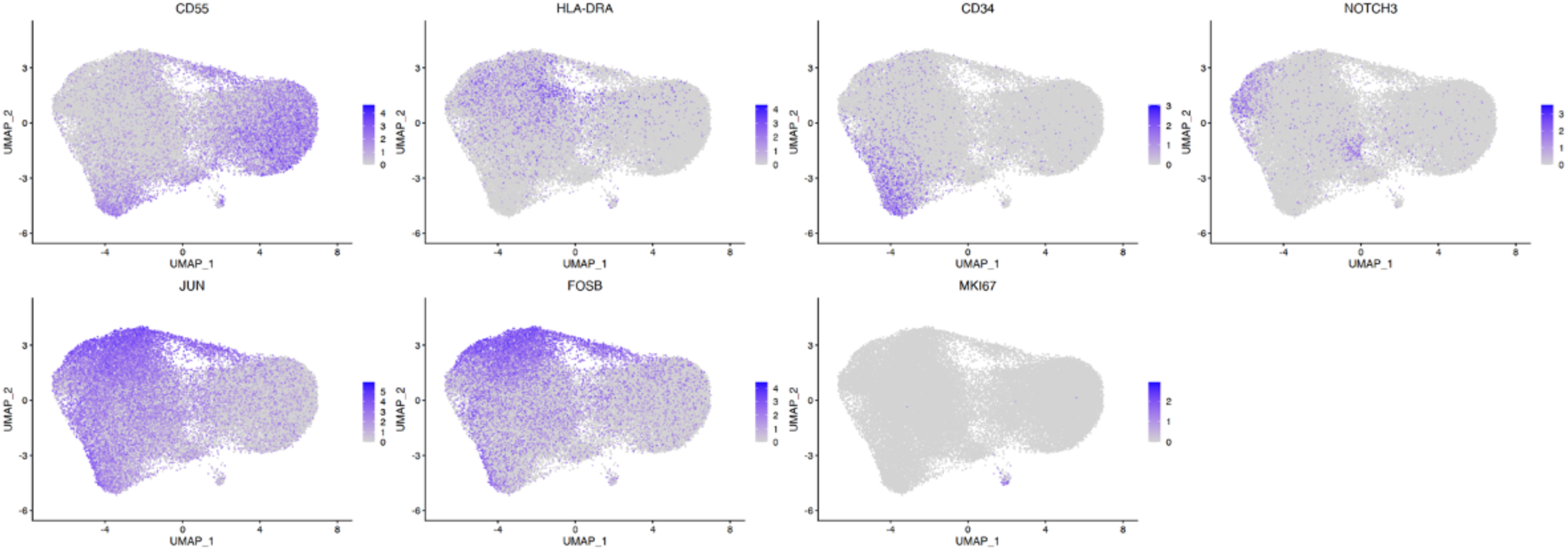
Expression of genes serving as markers of major fibroblast subsets in synovial fibroblasts. UMAP showing expression of the indicated gene from a dataset^11^ of 47,579 fibroblasts collected from synovial biopsies from OA (n=6) and RA (n=15).

**Extended Data Figure 3.**
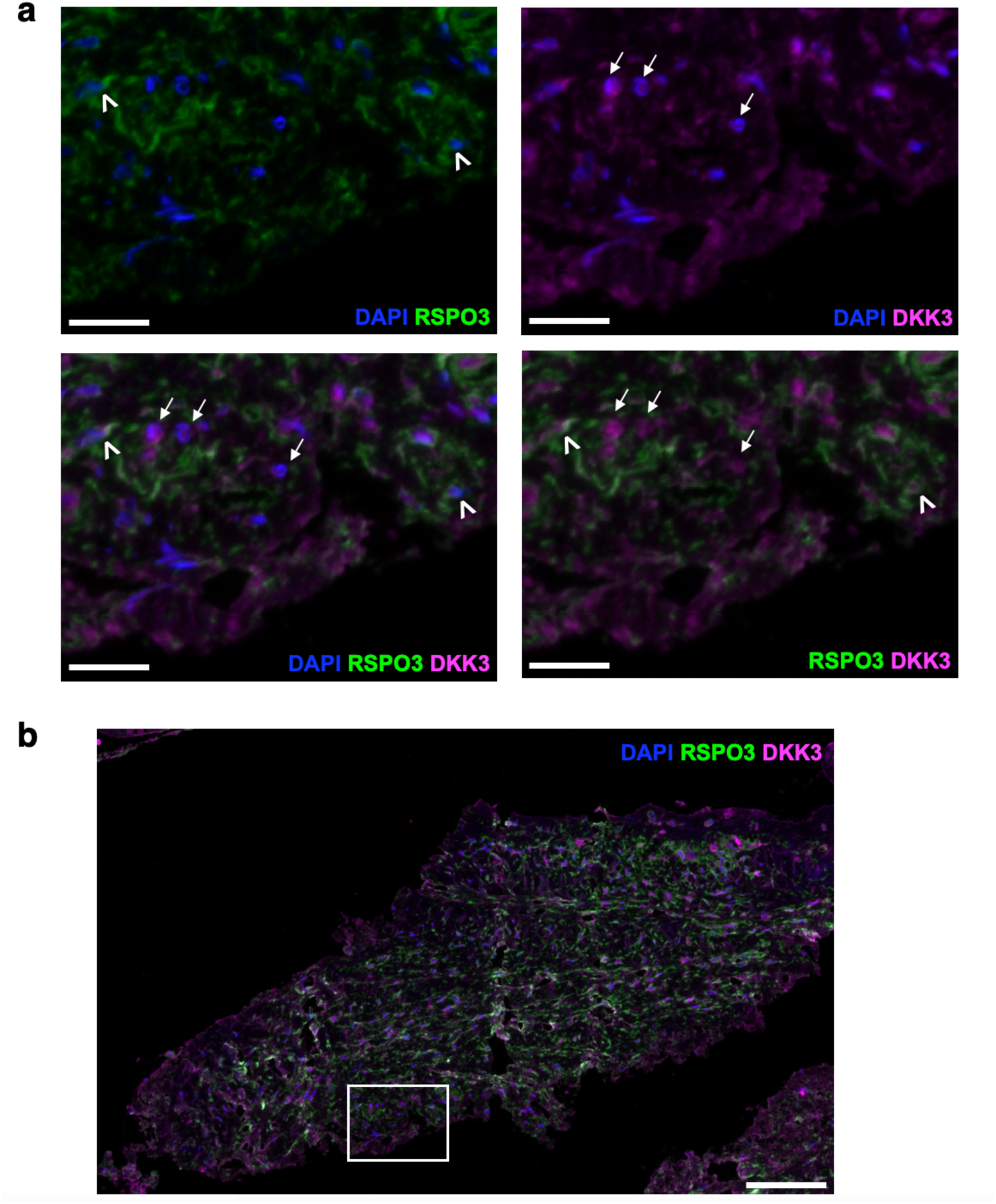
RSPO3 and DKK3 proteins show largely reciprocal patterns of expression. **a**, Multispectral immunofluorescence staining of RA synovial tissue highlighting expression of RSPO3 (green), DKK3 (magenta), and DAPI (blue). Carets indicate fibroblasts with high expression of RSPO3 but low expression of DKK3 and arrows indicate fibroblasts with high expression of DKK3 and low expression of RSPO3. Scale bar, 20 μM. **b**, Extended view of multispectral immunofluorescence staining of RA synovial tissue shown in **a**. Box indicates area depicted in **a**. Scale bar, 100 μM.

**Extended Data Figure 4.**
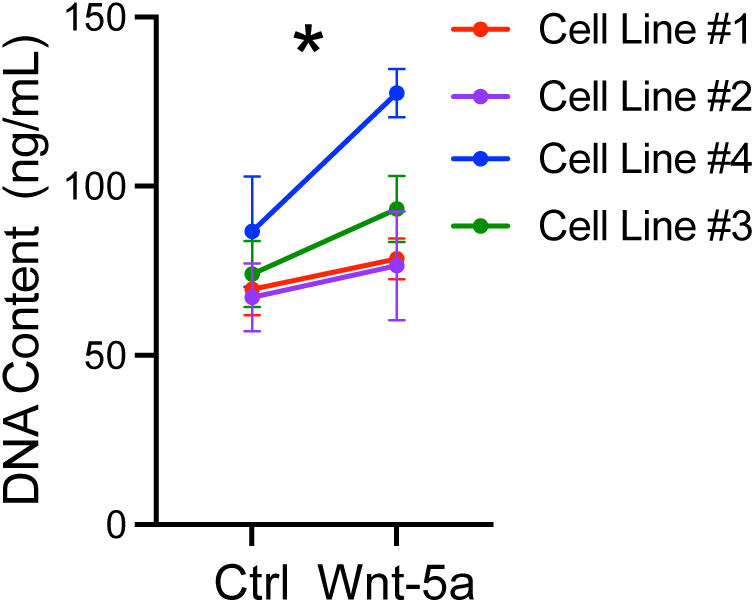
Non-canonical Wnt activation results in enhanced proliferation of RA synovial fibroblasts. Plot of DNA content from cultured RA synovial fibroblast cell lines 4 days post-stimulation with recombinant Wnt-5a or control. 4 cell lines are depicted with N=3 well replicates per cell line. Mean ± SD is shown. Normality was assessed using the Shapiro-Wilk test and significance was calculated using the paired Student’s T test of the ratio of Wnt-5a treated cells versus control. *p<0.05.

**Extended Data Figure 5.**
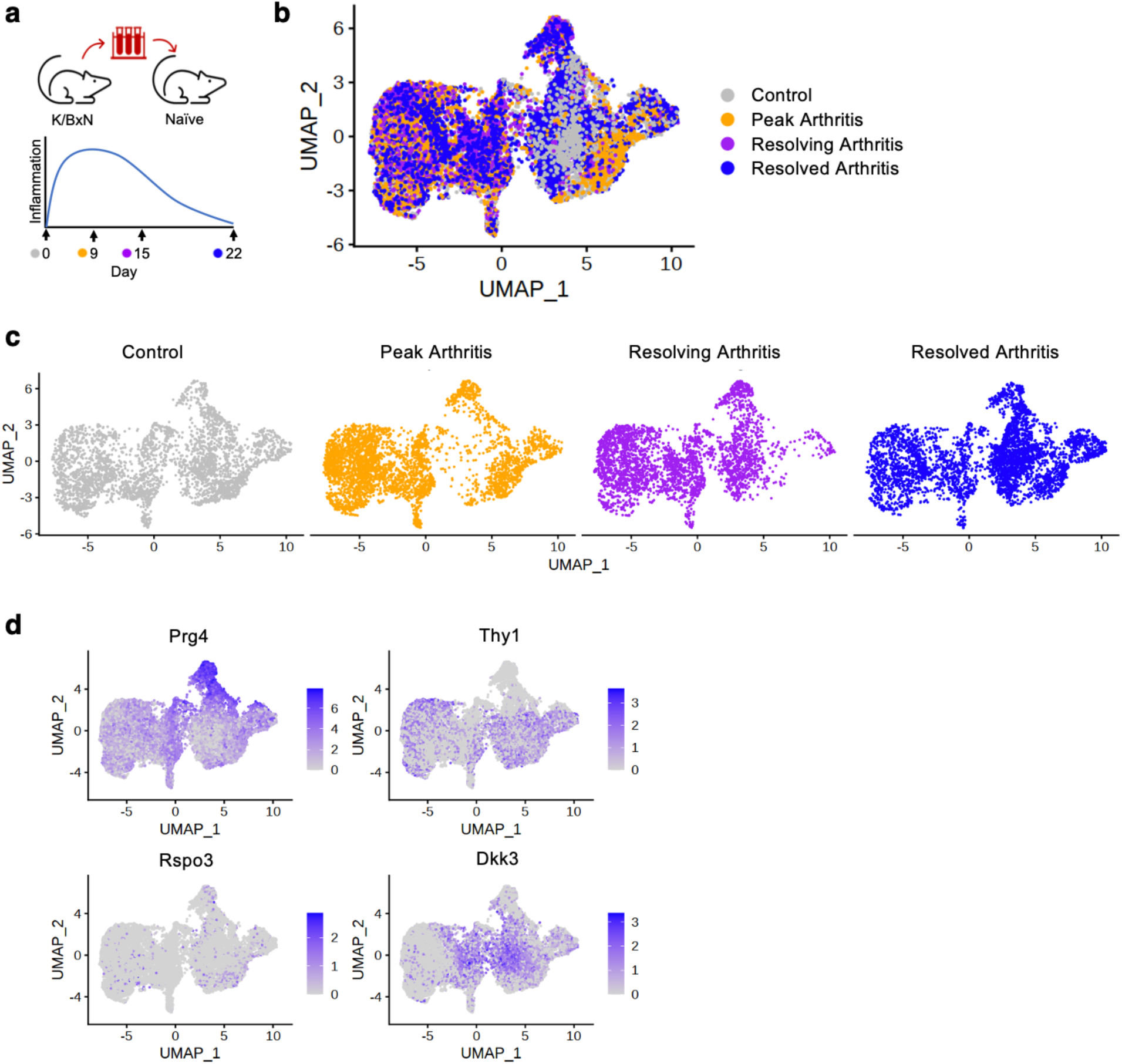
Wnt signaling is enriched at the peak of arthritis in a mouse model of inflammatory arthritis. **a**, Schematic depicting K/BxN arthritis model. **b**-**c**, UMAPs showing 14,541 synovial fibroblasts from K/BxN mice at varying time points in the arthritis course with time points overlaid (**b**) or separated (**c**). **d**, UMAP showing expression of the indicated markers in synovial fibroblasts from the dataset in **b**. For **b**-**d**, n=3 biologic replicates per time point.

**Extended Data Figure 6:**
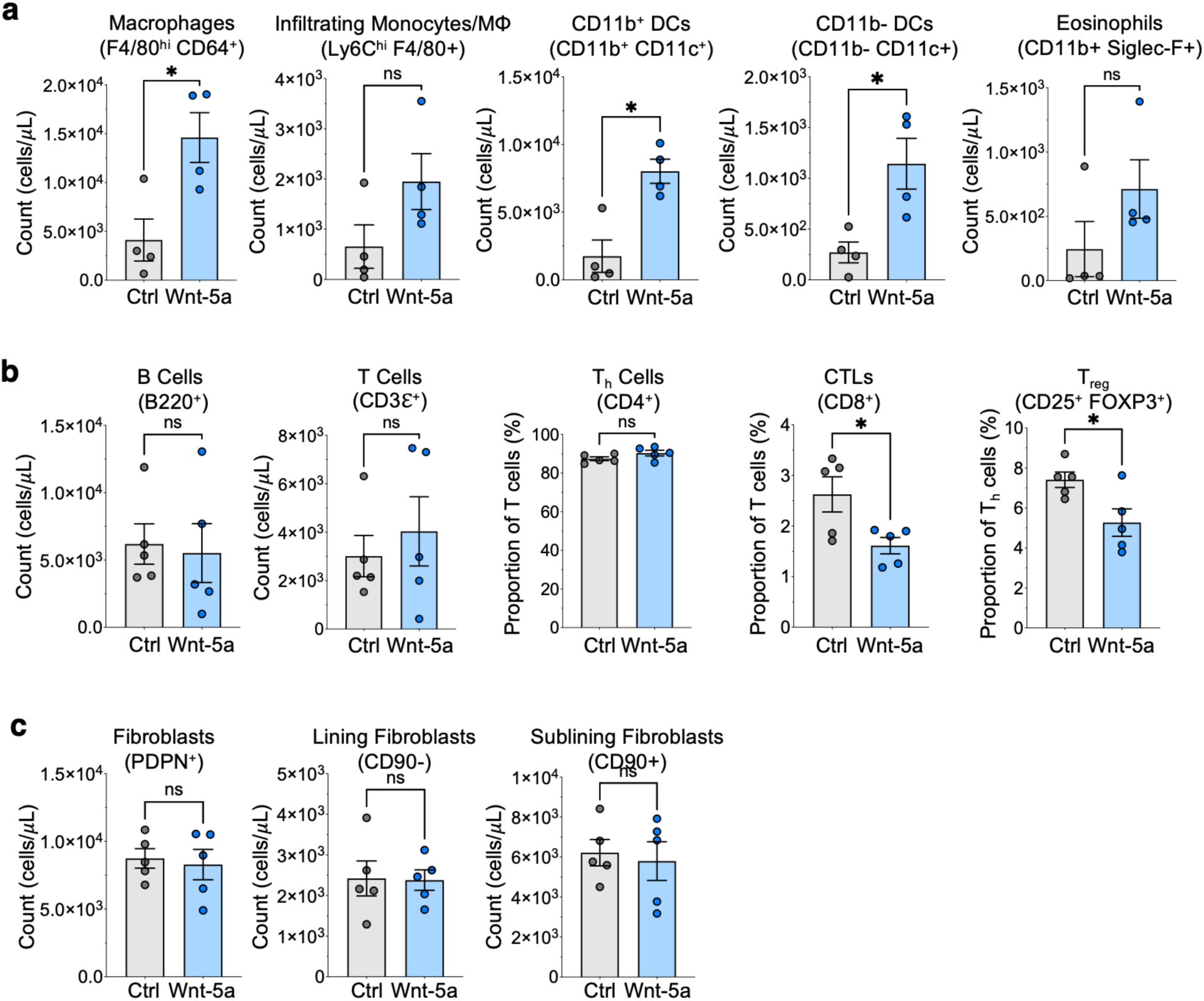
Wnt-5a activation results in an increase in dendritic cells, though no significant changes in T Cell, B Cell, eosinophil, or fibroblast numbers. **a-c**, Flow cytometry depicting absolute numbers of the indicated cell populations including myeloid cells (**a**), lymphocytes (**b**), and fibroblasts (**c**) isolated from knee synovial tissue from mice undergoing AIA treated with Wnt-5a or control. In **a**, n=5 per condition for plots of T cells and B cells and n=4 per condition for other plots. In **b** and **c**, n=5 per condition. For all plots, mean ± SEM is shown. The Shapiro-Wilk test was used to determine normality and p-values were calculated using a Student’s t-test for normally distributed data and Mann-Whitney test for non-normal distributions. *p<0.05.

**Extended Data Figure 7:**
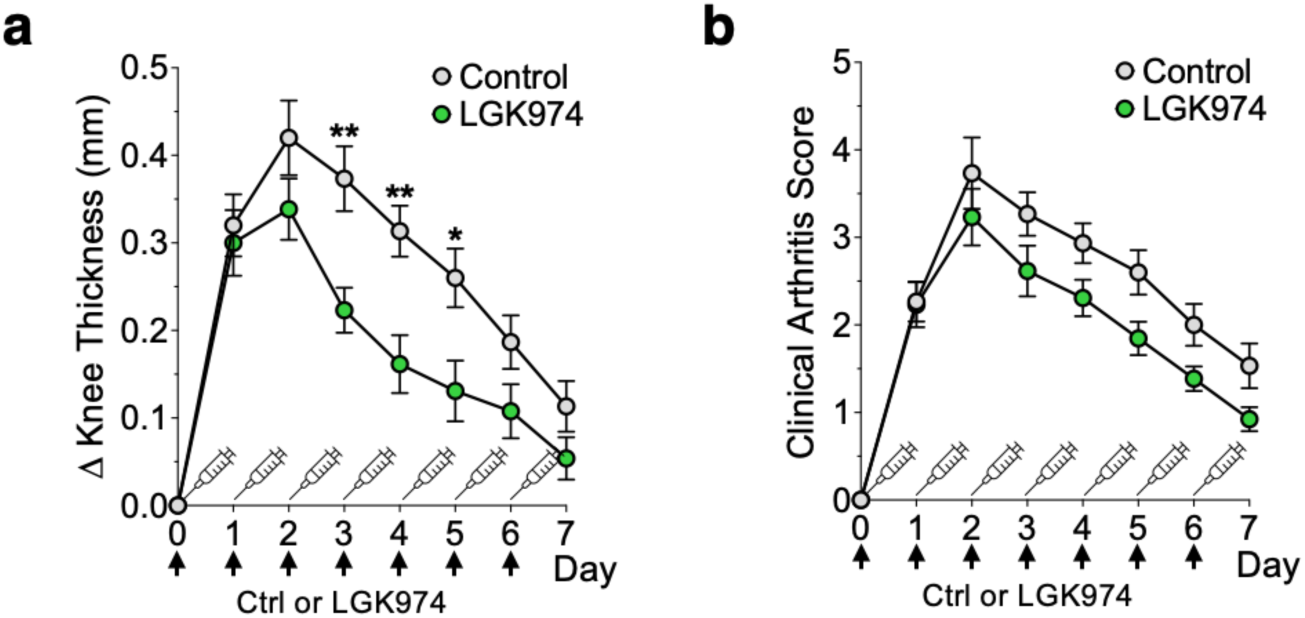
Treatment with LGK974 decreases arthritis severity. **a**, Knee thickness measurements of mice undergoing AIA that were treated with recombinant LGK974 or a vehicle control at the indicated time points, continuing to Day 7. **b**, Global arthritis scores for mice treated with LGK974 or vehicle control during AIA continuing to Day 7. For **a** and **b**, mean ± SEM comparing vehicle control (n=15) and LGK974 mice (n=13) is shown. Significance was calculated using a 2-way ANOVA with Šídák’s multiple comparisons test. *p<0.05, **p<0.01.

**Extended Data Figure 8:**
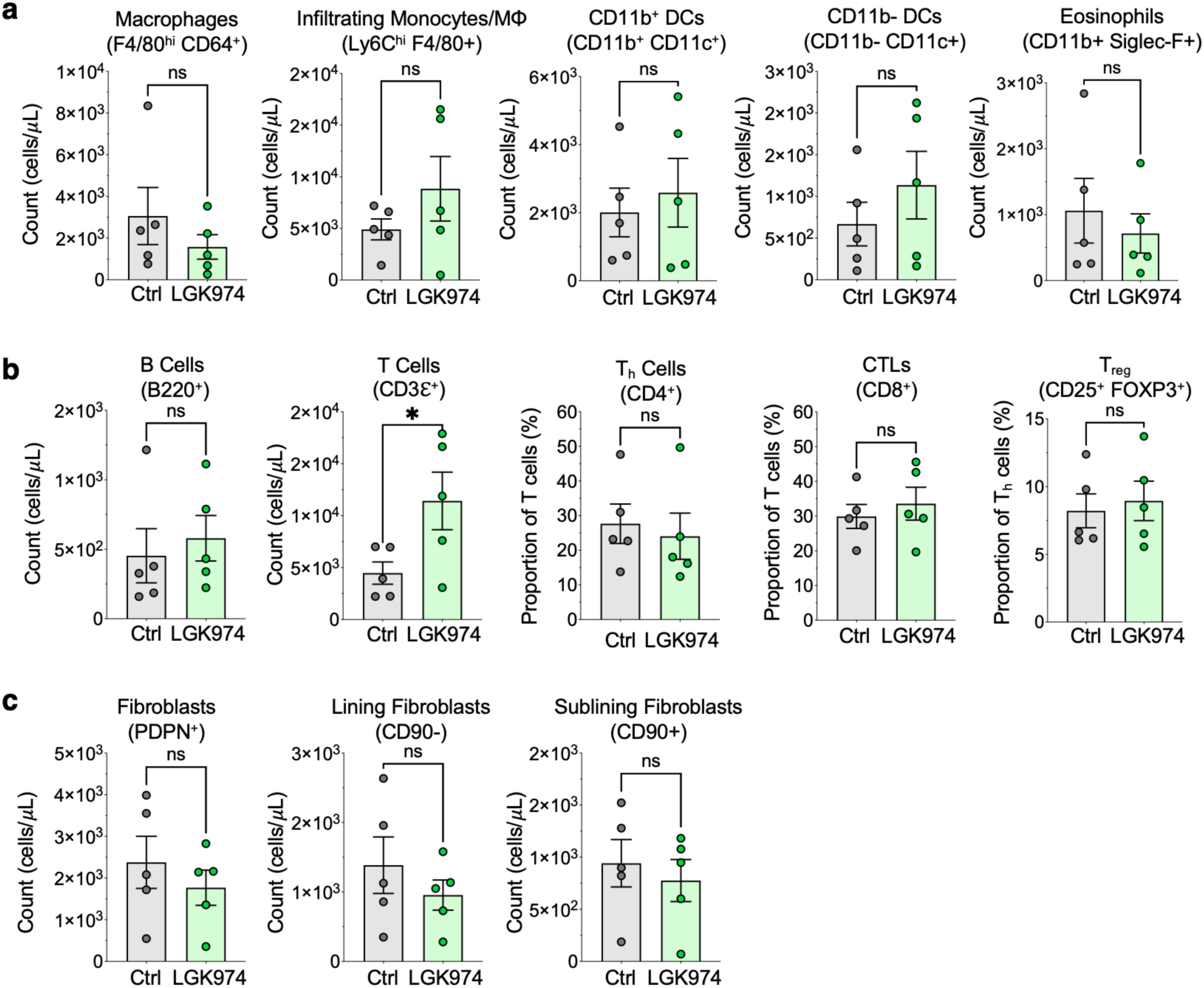
Wnt inhibition with LGK974 does not affect numbers of B cell, eosinophil, dendritic cell, monocyte, fibroblasts though there is an increase in T cell numbers. **a-c**, Flow cytometry depicting absolute numbers of the indicated cell populations including myeloid cells (**a**), lymphocytes (**b**), and fibroblasts (**c**) isolated from knee synovial tissue from mice undergoing AIA treated with LGK974 (n=5) or controls (n=5). For all plots, mean ± SEM is shown. The Shapiro-Wilk test was used to determine normality and p-values were calculated using a Student’s t-test for normally distributed data and Mann-Whitney test for non-normal distributions. *p<0.05.

## Methods

### Cell Culture

Synovial fibroblast cell lines used in this study were derived from synovial tissue of male and female adult patients with seropositive and seronegative RA per 2010 ACR/EULAR criteria. Samples were obtained from patients with active disease through synovectomies and biopsies or with chronic disease through arthroplasties. Samples used for genomic studies were obtained from patients consented through the following hospital IRB protocols: 2014P002558 and IRB 2019P002924. Cells were cultured at 37°C in 10% CO2 with the “FLS media” composed of the following ingredients: Dulbecco’s Modified Eagle Medium (DMEM) (Gibco, 11995-065), 1% MEM Non-Essential Amino Acids (Gibco, 11140-050), 2% MEM Amino Acids (Gibco, 11130-051), 1% Pen Strep (Gibco, 15140-122), 2mM L-Glutamine (Gibco, 25030-081), 55uM 2-Mercaptoethanol (Gibco, 21985-023), 10ug/mL Gentamicin (Gibco, 15710-064), 10 mM HEPES (Gibco, 15630-080), and 10% Fetal Bovine Serum (FBS) (Gemini Bio, 100-106).

### Bulk RNA sequencing

Fibroblasts were plated at 10,000 cells per well in FLS media in 96-well flat-bottom microplates (Fisher Scientific, 07-200-90) for 24 hours. The following day, the media was replaced with FLS starvation media supplemented with 1% FBS as described above. After 24 hours, cells were incubated with Wnt-3a (R&D, 5036-WN/CF) at 10 ng/mL, 50 ng/mL, or 500 ng/mL, Wnt-5a (R&D, 645-WN/CF) at 10 ng/mL, 50 ng/mL, or 1000 ng/mL or an equal volume of PBS (Gibco, 10010-023) in starvation media. After stimulation with Wnt ligand or controls for 4 hours or 16 hours, the media was removed, and the cells were rinsed with PBS. TCL buffer (Qiagen, 1031576) was then applied.

Sequencing libraries were prepared using the Illumina Smart-Seq2 protocol and sequenced on MiSeq from Illumina, generating 35 base paired-end reads. Reads were mapped to the Gencode Release 33 transcriptome using Kallisto 0.46.2^57^ to quantify transcripts per million (TPM) for all genes. Samples with fewer than 1 million reads were excluded from downstream analyses. To identify genes upregulated or downregulated with maximal doses of Wnt-3a and Wnt-5a at each time point, differential gene expression was assessed with DeSeq2^58^ where the tximport R package^59^ to import data from Kallisto. Second, to identify genes that showed a dose-responsive increase in expression with increased Wnt ligand dose, the limma R package^60^ was used. Here, linear models were developed to evaluate change in gene expression over log(ligand concentration) and the slope for each gene at each time point and Wnt ligand was assessed. The list of top 50 genes with the highest slope on linear modeling analysis or highest fold change on differential gene expression analysis were used for enrichment score calculations using a weighted scoring system from the singlecellmethods Github repository (https://github.com/immunogenomics/singlecellmethods) for analysis of bulk RNA sequencing data and human RA synovial scRNA-seq data or the Seurat addModuleScore function^61^ for analysis of murine scRNA-seq data.

### ELISA

Fibroblasts from the indicated cell lines were plated at 10,000 cells per well in 200 µL of FLS media in a 96-well flat-bottom tissue culture plate (Corning 3596) for 24 hours. Cells were stimulated with Wnt-3a (R&D 5036-WN/CF) at 500ng/mL, Wnt-5a (R&D 645-WN/CF) at 500ng/mL, or an equal volume of PBS for 16 hours. Supernatants were processed for ELISA measurements of IL-6 (R&D, DY206) and IL-8 (R&D, DY208) using a three-fold dilution.

### Cell Proliferation Assay

Fibroblasts were plated at 2000 cells per well in FLS media in 96-well optical-bottom plates (Thermo Fisher Scientific, 165305) for 24 hours. The following day, the media was replaced with 200 µL of FLS media supplemented with 1% FBS rather than 10% FBS (“starvation media”). After 24 hours, cells were then stimulated with either 500 ng/mL Wnt-3a (R&D, 5036-WN/CF), 500 ng/mL Wnt-5a (R&D, 645-WN/CF) or an equal volume of PBS (Gibco, 10010-023) in starvation media. Four days after stimulation, cells were rinsed in PBS and the plate was stored at −80°C. For measurement of DNA content, cells were lysed in buffer containing RNase A (Thermo Fisher Scientific, EN0531) using the standard protocol from the CyQUANT^TM^ Cell Proliferation Assay (Invitrogen, C7026).

### Human single-cell RNA-seq data processing and analysis

Count-level data and metadata are available from the following publicly available fibroblast single-cell RNA-seq datasets: AMP RA/SLE Phase 1 (ImmPort, accession SDY998 https://www.immport.org/shared/study/SDY998),^12^ Roche Fibroblast Network Consortium (Broad Single Cell Portal, accession SCP738; https://singlecell.broadinstitute.org/single_cell/study/SCP738/cross-tissue-single-cell-stromal-atlas-identifies-shared-pathological-fibroblast-phenotypes-in-four-chronic-inflammatory-diseases),^11^ and AMP RA/SLE Phase 2 (Synapse, accession 52297480; https://www.synapse.org/Synapse:syn52297840.7/datasets/).^7^ The Seurat R package (v4.3.0)^61^ for data processing and visualization. Each cell was normalized by multiplicatively scaling to 10,000 reads followed by log normalization (NormalizeData function). The top 2,000 most highly variable genes were selected by the VST method (FindVariableFeatures function). Highly variable genes were mean centered and unit variance scaled (scaleData function), and PCA was performed on these values with the top 30 PCs retained for subsequent analysis (runPCA function). The Harmony algorithm (v.1.0)^62^ was employed with default parameters to correct for donor-specific batch effects. For cell clustering, graph-based was performed using Louvain clustering (FindNeighbors and FindClusters functions) with resolution set to 0.3. Cluster marker genes were identified using the Wilcoxon rank sum test with Bonferroni p-value correction, excluding genes expressed by less than 20% of cells in either cluster or differentially expressed by less than an absolute log fold change of 0.5. Cells were visualized using UMAP projected onto two dimensions (runUMAP function). For the AMP Phase 1 dataset, cell projections into tSNE space from the original study were used for final visualization and gene expression plotting.

### Trajectory analysis

To identify genes that covary with RSPO3 and DKK3 in synovial fibroblasts, a trajectory was constructed from RSPO3-expressing fibroblasts to DKK3-expressing fibroblasts using the Roche Fibroblast Network Consortium synovial fibroblast dataset.^11^ In our preliminary analyses, we found that gene expression in this dataset was heavily influenced by donor-specific variability. To prevent donor effects from confounding the trajectory analysis, the donor-corrected cell embeddings generated by Harmony were used. In the Harmony embedding, we observed that RSPO3+ cells aggregated at one end while DKK3+ cells aggregated on the other, suggesting that a simple linear trajectory with no branching events would be sufficient to interrogate gene expression differences between RSPO3+ and DKK3+ fibroblasts. The trajectory was therefore designated as a line passing through the centroid of RSPO3+ cells and the centroid of DKK3+ cells. Pseudotime was assigned to cells based on their position along the linear trajectory using the princurve R package (v2.1.6).^63^

The limma package (v3.50.3)^60^ was then used to examine changes in gene expression across pseudotime. To model gene expression continuously over pseudotime, a regression spline was used with 5 degrees of freedom. Corrections were applied for gene expression variation attributable to donor or to fibroblast positional identity. Based on these factors, the following model was fit: GeneExpression ∼ PseudotimeSpline + DonorID + PositionalID + PositionalID:PseudotimeSpline. PseudotimeSpline represents the 5 parameters fit by the regression spline. DonorID indicates the donor from which each cell was derived. PositionalID indicates whether the cell was derived from the synovial lining, intermediate, or sublining region. PositionalID:PseudotimeSpline represents the interaction between positional identity and regression spline; we observed that a particular gene’s change over pseudotime could vary between fibroblasts derived from different regions of the synovium. Hierarchical clustering on the model coefficients for each gene was subsequently performed. Genes that co-clustered with RSPO3 and DKK3 were identified to covary with RSPO3 and DKK3.

### 6-plex multicolour immunofluorescence staining and image acquisition

RA synovial tissues were processed and stained using techniques previously described.^7^ Antibodies used for multiplex tissue immunofluorescence (IF) staining for RSPO3 (Clone 1E4F1, Proteintech, 1:100), VE-cadherin (Clone E6N7A, CST, 1:100) CD3 (Clone F7.2.38, DAKO, 1:200), Podoplanin/PDPN (EPR7072, Abcam, 1:100), CD90 (clone D3V8A, CST, 1:100), DKK3 (10365-1-AP, Proteintech, 1:200) and CD45 (clone D9M8I, CST, 1:100) was performed. Immunofluorescence signals for RSPO3, VE-cadherin, CD3, PDPN+CD90 (to characterize fibroblasts from both the lining layer and the sub-lining layer), DKK3, and CD45 were visualized using TSA dyes 570, 520, 620, 480, 690 and 780 respectively, and counterstained with spectral DAPI. Multispectral image processing of multiplex stained tissue and control tissue (which was stained with secondary only for DKK3 and RSPO3 markers) was performed using Phenochart (version 1.0.11/Akoya) and inForm Image Analysis Software (version 2.3, Akoya).

### Animal Husbandry

All animal experiments were approved by the UK Home Office and conducted in accordance with the UK’s Animals (Scientific Procedures) Act 1986 and the UK Home Office Code of Practice. The project and experimental protocols were approved by the University of Birmingham Animal Ethics Review Committee who provided ethical oversight of the study. C57BL/6 mice were purchased from Charles River (Strain 027). All mice were housed at a barrier and specific pathogen-free facility at the Biomedical Services Unit, University of Birmingham. All mice used in experimental studies were aged 6-10 weeks and unless otherwise stated, male. Single animals were considered as experimental units.

### K/BxN Serum Transfer Induced Arthritis

K/BxN serum transfer-induced arthritis (STIA) was induced by intravenous injection of 100µl of serum from KRN mice (Gifted from Harris Perlman, Department of Medicine, Division of Rheumatology, Northwestern University, Feinberg School of Medicine Chicago, Evanston, IL, USA). Inflammation of ankle and wrist joints was assessed daily, by measuring thickness, using callipers. Following induction of arthritis, tissues were harvested at days 0, 9,15 and 22 following induction of arthritis for scRNA-seq.

### Antigen-induced arthritis

For antigen-induced arthritis (AIA) mice were inoculated by injection of emulsified incomplete Freund’s adjuvant (Sigma Aldrich, F5506) supplemented with 2 mg/mL Mycobacterium tuberculosis H37Ra (BD, 231141), and 1mg/mL methylated-bovine serum albumin (m-BSA) (Sigma Aldrich, A1009), subcutaneously into either lateral side of the lower spine (50µL per injection site). 21 days following subcutaneous injections (Day 0), 100µg m-BSA in PBS was injected locally into the knee joint via intra-articular injection to induce arthritis. Inflammation of knee joints were assessed daily, by measuring thickness, using callipers.

### Injection of recombinant Wnt-5a in murine inflammatory arthritis model

6-10-week-old C57BL/6 mice were induced with AIA as previously described. At 48 and 96 hours following induction of inflammation (days 2 and 4, respectively), the knee was injected intra-articularly with 10µL recombinant Wnt-5a protein (R&D, 645-WN-010) diluted 1mg/mL in PBS or with a PBS control. The knee thickness was measured daily by calliper measurement of the knee joint. A global arthritis score was calculated adapted from an arthritis scoring system previously described.^64^ Following the induction of arthritis, tissues were harvested at day 7 for scRNA-seq, flow cytometry and histology.

### Injection of LKG974 in murine inflammatory arthritis model

6-10-week-old C57BL/6 mice were induced with AIA as previously described. LGK974 (3mg/kg Cambridge Biosciences, L2540) dissolved in corn oil (Sigma Aldrich, C8267) + 10% DMSO (Sigma Aldrich, D4540) or a vehicle control of corn oil (Sigma Aldrich, C8267) + 10% DMSO (Sigma Aldrich, D4540) was administered via intraperitoneal injection daily for 7 days (days 0-6). Knee thickness and global arthritis scores were assessed daily as described above. Following induction of arthritis, tissues were harvested at day 4 for scRNA-seq, flow cytometry and histology.

### Histology of murine synovial tissue

The tissue was fixed for 24 hours in 10% formalin, then decalcified in EDTA (10% w/v, pH 7.4). The tissue was then processed with the following protocol: 70% ethanol for 1 hour, 95% ethanol for 1 hour, 95% ethanol for 1 hour, 100% ethanol for 1 hour, 100% ethanol for 1 hour, and then incubation in 100% ethanol overnight. Sections were then placed in xylene for 1 hour. Fresh xylene was then applied for an additional hour. The tissue was then placed in paraffin (60°C) for 4 hours with the paraffin solution changed every 15 minutes. Cells were then embedded in paraffin and left to set at room temperature. 4µM sections were cut and incubated at 56°C overnight, then stored at 4°C. Sections were stained with H&E using standard procedures. Sections were imaged using the AxioScan Slide Scanner and analysed using ZenBlue software. Synovitis severity was determined using an adapted scoring system from published works.^10,65^ The sum of the individual scores for synovial hyperplasia (0-3), cellularity (0-3), and inflammation (0-3) was calculated to obtain a total synovitis score (0-9).

### Isolation of murine synovial cells

For analyses of fibroblast and myeloid cell fractions, the collagenase D/Dispase method was used. Briefly, bones with intact joints were dissected and transferred into RPMI-1640 (21870084, Gibco) +2% FCS (Biosera, FB-1001) containing 0.1 g/mL collagenase D (Roche, 11088866001), and 0.01 g/mL of DNase I (Sigma-Aldrich, DN25-10MG). Samples were incubated at 37 °C for 45 min, followed by incubation with medium containing 0.1 g/mL Collagenase/Dispase (Roche, 11097113001) and 0.01 g/mL DNase I at 37 °C for 30 min. The cell suspension was filtered through 50µm cell strainer (Corning, CLS431751), and red blood cell lysis (Sigma Aldrich, R7757) was performed. The resulting cells were pelleted by centrifugation at 400g for 5 minutes.

For analyses of lymphocytes, the Collagenase P method was applied. Bones with intact joints were dissected and transferred into RPMI-1640 (+2% FCS) containing 0.8 g/mL Collagenase/Dispase, 0.2 mg/mL collagenase P (Roche, 11213857001) and 0.1 mg/mL of DNase I. Samples were incubated at 37 °C for 30 minutes, followed a second incubation with fresh digest medium for 20 minutes. The cell suspension was filtered through 50µm cell strainer, and red blood cell lysis was performed. The resulting cells were pelleted by centrifugation at 400g for 5 minutes.

### Flow cytometry of murine synovial cells

Cells were harvested using the digest protocol specified as above and pelleted (400g, 5 minutes, 4°C). For cell staining, cells were washed twice with PBS before incubating with fixable viability dye (Biolegend, 423101). Antibody cocktails were prepared using FACS buffer (PBS + 0.1% BSA). Cells were washed twice with FACS buffer and stained with antibody mix (Tables 1–4). Cells were washed twice with FACS buffer and prepared for analysis. For Foxp3 staining following staining of cell surface markers, cells were fixed/permeabilised using Foxp3 / Transcription Factor Staining Buffer Set (Invitrogen, 00-5523-00). Cells were washed twice with 1X permeabilisation buffer, and incubated with mouse FC block (Biolegend, 156603), then α-FOXP3 was added (Invitrogen, 12-5773-82). Cells were washed twice with FACS buffer and prepared for analysis. To assess fibroblast activation, following fixable viability staining cells were blocked then incubated with α-FAPα (BioTechne, AF3715). Cells were washed twice in FACS buffer, blocked and incubated with biotinylated α-sheep (Sigma Aldrich, B3148). Cells were then washed twice in FACS buffer and incubated in the antibody cocktail containing all extracellular markers. Once staining was completed, the volume of each sample was measured before addition of counting beads (Invitrogen, C36995). Samples were run on a BD LSRFortessa X-20 and analysed using FlowJo v10.

**Table 1:** Fibroblast Antibody Panel.

| **Target** | **Fluorochrome** | **Clone** | **Supplier** | **Catalogue #** | **Dilution** |
| --- | --- | --- | --- | --- | --- |
| CD45 | PE-CY7 | 30-F11 | Biolegend | 103113 | 1/500 |
| CD31 | SuperBright-645 | 390 | eBioscience | 64-0311-82 | 1/500 |
| PDPN | PE | 8.1.1 | Biolegend | 127407 | 1/150 |
| CD90.2 | APC-CY7 | 30-H12 | Biolegend | 105327 | 1/100 |
| FAPα | - | Polyclonal | BioTechne | AF3715 | 1/50 |
| Biotinylated α-sheep | - | GT-34 | Sigma Aldrich | B3148 | 1/1000 |
| Streptavidin | APC | - | Biolegend | 405207 | 1/400 |
| TruStain FcX<br>(anti-mouse<br>CD16/32) | - | S17011E | Biolegend | 156603 | 1/100 |
| Fixable<br>Viability | BV510 | - | Biolegend | 423101 | 1/500 |

**Table 2:** Macrophage Antibody Panel.

| **Target** | **Fluorochrome** | **Clone** | **Supplier** | **Catalog #** | **Dilution** |
| --- | --- | --- | --- | --- | --- |
| CD45 | APC-CY7 | 30-F11 | Biolegend | 103116 | 1/500 |
| CD11b | FITC | M1/70 | Biolegend | 101206 | 1/100 |
| LY6G | AF700 | 1A8 | Biolegend | 127622 | 1/100 |
| SIGLEC-F | PE-e610 | 1RNM44N | Invitrogen | 46-1702-2 | 1/200 |
| CD64 | PE-CY7 | X54-5/7.1 | Biolegend | 139314 | 1/100 |
| F4/80 | BV421 | BM8 | Biolegend | 123131 | 1/100 |
| CX<sub>3</sub>CR1 | PerCP-Cy5.5 | SA011F11 | Biolegend | 149010 | 1/100 |
| MHC II | BV650 | M5/114.15.2 | Biolegend | 107641 | 1/200 |
| LYVE1 | PE | ALY7 | Invitrogen | 12-0443-82 | 1/100 |
| MerTK | APC | 2B10C42 | Biolegend | 151508 | 1/100 |
| Fixable<br>Viability | BV510 | - | Biolegend | 423101 | 1/500 |

**Table 3:**
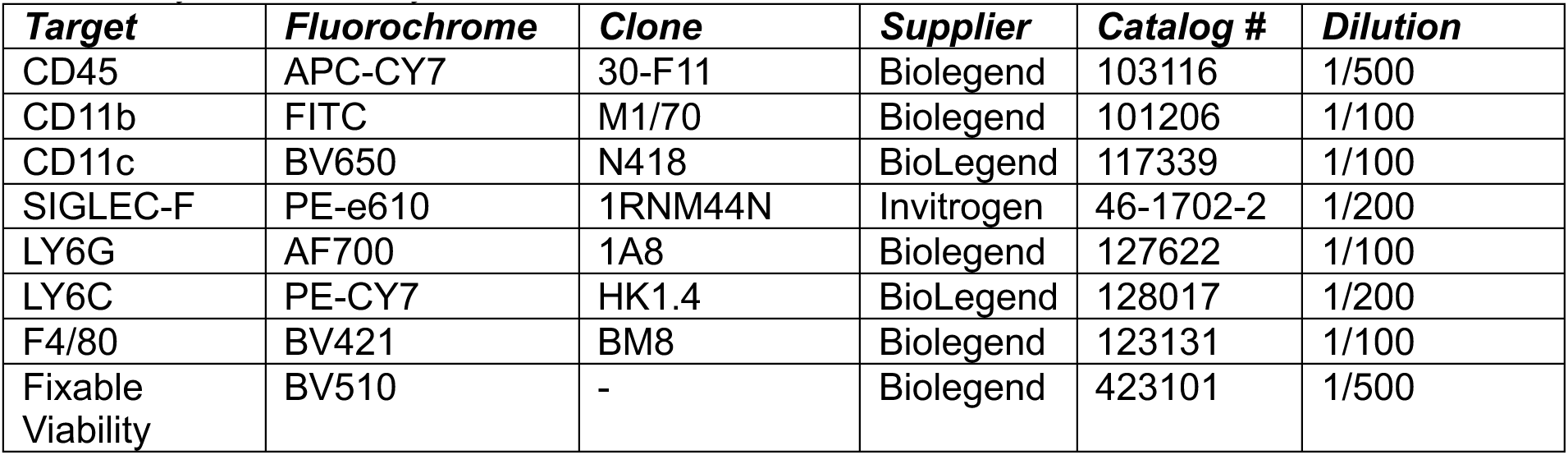
Myeloid Antibody Panel.

**Table 4:** Lymphocyte antibody panel.

| **Target** | **Fluorochrome** | **Clone** | **Supplier** | **Catalog #** | **Dilution** |
| --- | --- | --- | --- | --- | --- |
| CD45 | PE-CY7 | 30-F11 | Biolegend | 103113 | 1/300 |
| CD8a | PE-Texas Red | 53-6.7 | Biolegend | 100761 | 1/100 |
| CD3e | APC-e780 | 145-2C11 | Biolegend | 100329 | 1/80 |
| B220 | FITC | RA3-6B2 | Biolegend | 103206 | 1/100 |
| CD4 | BV421 | RM4-5 | Biolegend | 100543 | 1/80 |
| CD25 | BV650 | PC61 | BioLegend | 102037 | 1/80 |
| FOXP3 | PE | FJK-16s | Invitrogen | 12-5773-82 | 1/50 |

### Single cell RNA sequencing of murine synovial samples

Isolated synovial cells were stained at 4°C and dead cells excluded using 7-Aminoactinomycin D (7-AAD) staining (ThermoFisher, A1310). Antibodies used were anti-CD45 (Biolegend, 30-F11). Cell sorting was performed immediately after staining using a MoFlo Astrios EQ machine (Beckman Coulter). Cells were captured with the 10X Genomics Chromium system, and sequencing libraries were generated using the 10X Genomics Single Cell 3′ Solution (version 2) kit then sequenced with Illumina sequencing (HiSeq 4000, read 2 sequenced to 75 bp). Alignment, quantitation and aggregation of sample count matrices was performed using the 10X Genomics Cell Ranger pipeline (v7.0.1) against the mouse transcriptome (10x Genomics 2024-A reference, GRCm39 assembly). Downstream analysis was performed using the Seurat R package (v.4.1.1).

